# Stage-Dependent Differential Gene Expression Profiles of Cranial Neural Crest Cells Derived from Mouse Induced Pluripotent Stem Cells

**DOI:** 10.1101/432302

**Authors:** Ayano Odashima, Shoko Onodera, Akiko Saito, Takashi Nakamura, Yuuki Ogihara, Tatsuya Ichinohe, Toshifumi Azuma

## Abstract

Cranial neural crest cells (cNCCs) comprise a multipotent population of cells that migrate into the pharyngeal arches of the vertebrate embryo and differentiate into a broad range of derivatives of the craniofacial organs. Consequently, migrating cNCCs are considered as one of the most attractive candidate sources of cells for regenerative medicine. In this study, we analyzed the gene expression profiles of cNCCs at different time points after induction by conducting three independent RNA sequencing experiments. We successfully induced cNCC formation from mouse induced pluripotent stem (miPS) cells by culturing them in neural crest inducing media for 14 days. We found that these cNCCs expressed several neural crest specifier genes but were lacking some previously reported specifiers, such as paired box 3 (*Pax3*), msh homeobox 1 (*Msx1*), and Forkhead box D3 (*FoxD3*), which are presumed to be essential for neural crest development in the embryo. Thus, a distinct molecular network may the control gene expression in miPS-derived cNCCs. We also found that *c-Myc*, ETS proto-oncogene 1, transcription factor (*Ets1*), and sex determining region Y-box 10 (*Sox10*) were only detected at 14 days after induction. Therefore, we assume that these genes would be useful markers for migratory cNCCs induced from miPS cells. Eventually, these cNCCs comprised a broad spectrum of protocadherin (Pcdh) and a disintegrin and metalloproteinase with thrombospondin motifs (Adamts) family proteins, which may be crucial in their migration.

## Introduction

Stem cell-based tissue engineering is important in the field of oral science as it allows the regeneration of damaged tissues or organs [1,2]. Various stem cell populations have been identified as having a regeneration potential in the craniofacial region; however, the cranial neural crest cells (cNCCs) are considered as one of the most important candidates due to their role in craniofacial tissue organization [3].

cNCCs comprise a multipotent population of migratory cells that are unique to the vertebrate embryo and give rise to a broad range of derivatives [4,5], with the neural crest (NC) being capable of forming teratoma when transplanted into the immunocompromised animals [6]. The development of cNCCs involves three stages [7–10]: the neural plate border stage, the premigratory stage, and the migratory stage. During the migratory stage, the cNCCs delaminate from the posterior midbrain and individual rhombomeres in the hindbrain [11] and migrate into the pharyngeal arches to form skeletal elements of the face and teeth and contribute to the pharyngeal glands (thymus, thyroid, and parathyroid) [12]. Consequently, presumably cNCCs may represent a new treatment strategy for diseases in the craniofacial region [13].

Development from the premigratory to migratory stage proceeds swiftly [14], making it difficult to isolate and characterize a pure cNCC population from the embryo [15]. A recent transcriptome analysis of pure populations of sex determining region Y-box 10 (*Sox10*) + migratory cNCCs from chicks [16] has greatly improved our understanding of the characteristics of cNCCs, and methods for deriving NCCs from the embryonic stem (ES) cells have also been reported [17–30]; however, it remains unclear whether these cells are in the migratory stage and how long it takes to promote ES cell-derived NCCs from the pre-migratory to migratory stage.

In recent years, the use of induced pluripotent stem (iPS) cells as a revolutionary approach to the treatment of various medical conditions has gained immense attention [31,32] and iPS cells have several clear advantages over ES cells and primary cultured cNCCs as a cell source in regenerative medicine [16]. NCCs have been generated from iPS cells in numerous ways [24,33–38], with two reports having examined the differentiation of NCCs from ES or iPS cells [24,39] and two articles having described the protocol for differentiating NCCs from mouse iPS (miPS) cells [33,34]; however, few studies have investigated the changes in the properties of these NCCs overtime during the dynamic differentiation processes in the NC, in particular, during the migratory stage. Embryonic NC development depends on several environmental factors that influence the NC progenitors, regulation, and the timing of differentiation, making the elucidation of the gene regulatory network and expression profiles of miPS cell-derived cNCCs important.

Recent advances in the next-generation RNA sequencing technology (RNA-seq) have made it possible to analyze the gene expression profiles comprehensively [40–42]. Therefore, here, we used RNA-seq to investigate the gene expression landscape of cNCCs induced from miPS cells.

We treated the iPS-derived cells with cNCC induction medium for 14 days and performed triplicate RNA-seq experiments. We found that standard NC markers such as nerve growth factor receptor (*Ngfr*), snail family transcriptional repressor 1 (*Snai1*), and *Snai2* were remarkably increased at 7 days after cNCC induction; whereas, the expression of the cNCC markers ETS proto-oncogene 1, transcription factor (*Ets1*), and *Sox5, -8, -9*, and *-10* characteristically increased at 14 days after cNCC induction. Nestin (*Nes*) was upregulated throughout cNCC differentiation, as described previously [23]. In contrast, the homeobox genes such as msh homeobox 1 (*Msx1*), paired box 3 (*Pax3*), and *Pax7* were not detected in the NC after a longer period of differentiation, despite their expressions having been observed in several animals [43–52]. Furthermore, the expression of Forkhead box D3 (*FoxD3*), which is known to be required for maintaining pluripotency in mouse ES cells [53] and is also an important NC specifier transcription factor during embryonic development, decreased over time, suggesting that it is not a cNCC specifier in iPS-derived cells.

Another important finding was the remarkable upregulation of several metzincins, including members of the disintegrin and metalloproteinase domain metallopeptidase with thrombospondin motifs (Adamts) metalloproteinase family, which play crucial roles in modulating the extracellular matrix (ECM) during development [54–56]. We assume that various kinds of Adamts proteins produce distinct extracellular proteins that are digested by cNCC swallowing them to easily migrate toward their final destinations. We also found that the expressions of nearly all procadherin (Pcdh) superfamily members were increased, some only at the migratory stage. Pcdh is the largest subfamily of cadherins and the digestion of Pcdh protein by Adam proteins is crucial for development [57].

Eventually, our results indicated that *c-Myc*; *Ets1*; *Sox10; Adamts2* and -*8*; protocadherin alpha 2 (*Pcdha2*)*; Pcdha5*, -7, *-11*, and *-12;* protocadherin alpha subfamily C,1 (*Pcdhac1*); and protocadherin gamma subfamily C, 3 (*Pcdhgc3*) may represent appropriate markers for migratory cNCCs induced from miPS cells.

## Materials and Methods

### miPS cell culture

All of the mouse studies were conducted in accordance with protocols approved by the Animal Research Committee of Tokyo Dental College (No. 270401).

The miPS cells that were used in this study (APS0001; iPS-MEF-Ng-20D-17 mouse induced pluripotent stem cell line) were purchased from RIKEN BRC (Ibaraki, Japan) [58]. The cells were maintained on inactivated murine embryonic fibroblast (MEF) feeder cells in Dulbecco’s Modified Eagle’s Medium (DMEM; Invitrogen, Carlsbad, CA, USA)supplemented with 15% KnockOut™ Serum Replacement (Invitrogen), 1% nonessential amino acids (Chemicon, Temecula, CA, USA), 1% L-glutamine (Chemicon), 1000 U/ml penicillin–streptomycin (P/S; Invitrogen), and 0.11 mM 2-mercaptoethanol (Wako Pure Chemical Industries Ltd., Osaka, Japan) and were passaged in 60-mm cell culture plates at a density of 1 × 10^5^ cells/plate. The cells were grown in 5% CO_2_ at 95% humidity and the culture medium was changed each day.

### Embryoid body (EB) formation and cNCC differentiation

We obtained cultured cNCC cells following a previously described procedure [59], as outlined in Fig 1. miPS cells were dissociated with 0.05% trypsin–ethylenediaminetetraacetic acid (EDTA; Invitrogen) and were transferred to low-attachment, 10-mm petri dishes at a density of 2 × 10^6^ cells/plate to generate EBs. The EBs were then cultured in the NC induction medium comprising a 1:1 mixture of DMEM and F12 nutrient mixture (Invitrogen) and Neurobasal™ medium (Invitrogen) supplemented with 0.5 × N2 (Invitrogen), 0.5 × B27 (Invitrogen), 20 ng/ml basic fibroblast growth factor (Reprocell, Yokohama, Japan), 20 ng/ml epidermal growth factor (Peprotech, Offenbach, Germany), and 1% penicillin–streptomycin (P/S) for 4 days, during which time the medium was changed every other day. After 4 days, the day 0 (d0) EBs were collected and plated on 60-mm cell culture plates coated with 1μg/ml collagen type I (Advanced BioMatrix, San Diego, CA, USA). The cells were then subcultured in the same medium, which was changed every other day, and any rosetta-forming cells were eliminated. After 7–10 days, d7 cells were dissociated with 0.05% trypsin–EDTA and transferred to 60-mm cell culture plates coated with 1μg/ml collagen type I at a density of 1 × 10^5^ cells/plate to generate 14 cells. The cells from each of these passages were collected for RNA extraction.

**Fig 1.**
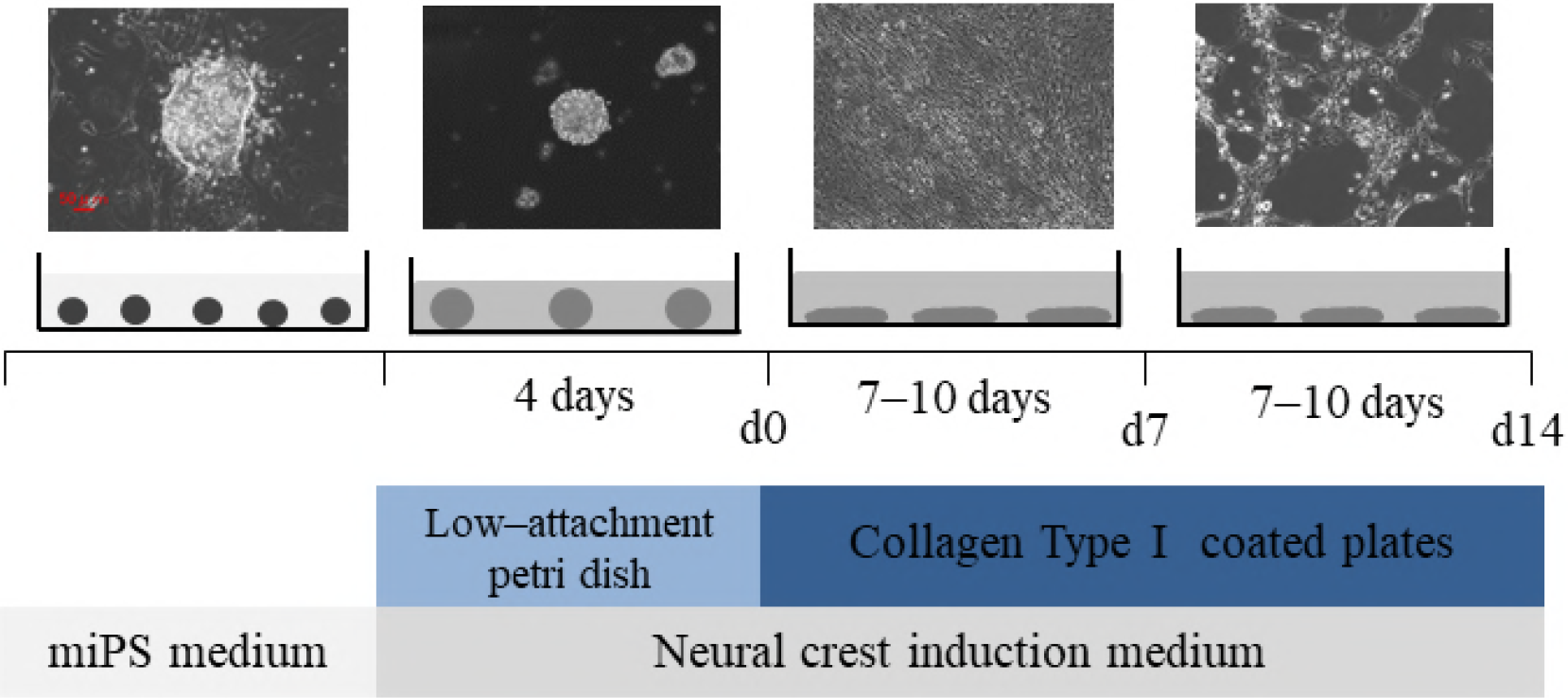
The experimental protocol that was used to induce the formation of cranial neural crest cells (cNCCs) from mouse induced pluripotent stem (miPS) cells. The photographs reveal miPS cells at four different stages: initial miPS cells, embryoid body (EB) on day 0 (d0), and cNCCs on d7 and d14. Small circles represent miPS cells; large circles represent EBs; ellipses represent d7 and d14 cells. Scale bar = 50 μm.

### O9–1 cell culture

O9–1 cells, which area mouse cNCC line, were purchased from Milliopore (Billerica, MA, USA) and cultured as previously described [50] as a control.

### RNA isolation and quantitative reverse transcription polymerase chain reaction analysis

The representative NC markers *Ngfr, Snai1, Snai2, Sox9*, and *Sox10* were selected and analyzed by quantitative reverse transcription polymerase chain reaction (qRT-PCR) analysis. Total RNA was extracted using QIAzol^®^ reagent (Qiagen, Valencia, CA, USA) according to the manufacturer’s protocol and RNA purity was assessed using a NanoDrop^®^ ND-1000 spectrophotometer (Thermo Fisher Scientific, Waltham, MA, USA), which revealed that each RNA sample had an A260/A280 ratio of >1.9. Complementary DNA (cDNA) was synthesized using a high-capacity cDNA reverse transcription kit (Applied Biosystems, Foster City, CA, USA) and qRT-PCR analysis was performed using Premix Ex Taq™ reagent (Takara Bio Inc., Otsu, Japan) according to the manufacturer’s protocol and the Applied Biosystems^®^ 7500 Fast Real-Time PCR System, with the primer sequences presented in Table 1. All samples were normalized to levels of 18S ribosomal RNA (*18S rRNA*). The relative expressions of the genes of interest were analyzed using the ΔΔ*Ct* method and were compared among the groups using analysis of variance (ANOVA) followed by the Bonferroni test where the significant differences were detected among the groups. A significance level of *p* < 0.05 was used for all analyses and all data are expressed as means ± standard deviations (SD).

**Table 1.**
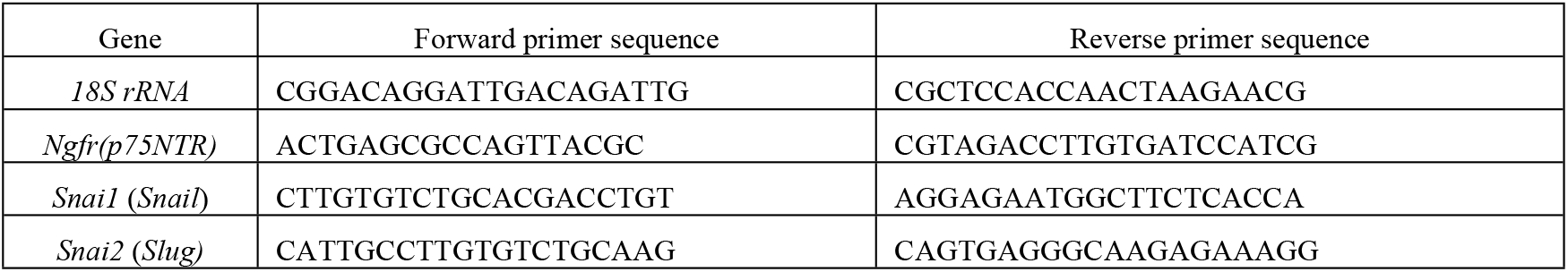
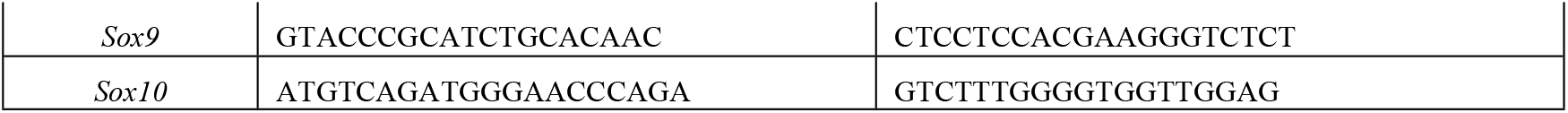
Primers used for quantitative reverse transcription polymerase chain reaction (qRT-PCR).

### Immunohistochemistry

The cells were fixed with 4% paraformaldehyde (Wako Pure Chemical Industries Ltd.) for 15 min followed by methanol (Wako Pure Chemical Industries Ltd) for 5 min. After washing, the nonspecific binding of antibodies was blocked by adding 5% bovine serum albumin (BSA; Wako Pure Chemical Industries Ltd.) in a phosphate buffered saline (PBS) with 0.5% Triton X-100 (PBST) for 1 h. The cells were then incubated with the primary antibodies Snai1 1:50 for goat anti-rabbit (Proteintech Group, Inc. Chicago, Il, USA) and Sox10 1:500 for goat anti-mouse (Atlas Antibodies, Bromma, Sweden) in PBST for 2 nights at 4 °C. They were then incubated in the secondary antibodies fluorenscein isothiocyanate conjugated anti-rabbit IgG (Abcam, Cambridge, MA, USA) at a dilution of 1:500 for Snai1 and anti-mouse IgG (Invitrogen) at a dilution of 1:500 for Sox10 in PBST for 1 h. Eventually, the cells were stained with 4,6-diamidino-2-phenylindole (DAPI; Sigma, Livonia, MI, USA) to visualize the nuclear DNA.

### RNA-seq and analysis

Total RNA from each sample was used to construct libraries with the Illumina TruSeq Stranded mRNA LT Sample Prep Kit (Illumina, San Diego, CA, USA), according to the manufacturer’s instructions. Polyadenylated mRNAs are commonly extracted using oligo-dT beads, following which the RNA is often fragmented to generate reads that cover the entire length of the transcripts. The standard Illumina approach relies on randomly primed double-stranded cDNA synthesis followed by end-repair, ligation of dsDNA adapters, and PCR amplification. The multiplexed libraries were sequenced as 125-bp paired-end reads using the Illumina Hiseq2500 system (Illumina). Prior to performing any analysis, we confirmed the quality of the data and undertook read cleaning, such as adapter removal and simple quality filtering, using Trimmomatic (ver. 0.32). The paired-end reads were then mapped to the mouse genome reference sequence GRCm38 using the Burrows–Wheeler Aligner (ver. 0.7.10). The number of sequence reads that were mapped to each gene domain using SAM tools (ver. 0.1.19) was counted and the reads per kilobase of transcript per 1 million mapped reads (RPKM) for known transcripts were calculated to normalize the expression level data to gene length and library size, allowing different samples to be compared.

## Results

### Gene expression profiles and immunohistochemistry of cNCCs differentiated from miPS cells

The expressions of the NC markers *Ngfr, Snai1, Snai2, Sox9*, and *Sox10* were examined by qRT-PCR in cNCCs differentiated from miPS cells as well as in O9–1 cells as a control. We detected the expression of all genes except *Ngfr* and *Sox10* in the O9–1 cells [50]. In contrast, all five genes were detected in the cNCCs, with the premigratory neural crest markers *Ngfr, Snai1*, and *Snai2* having the highest expression levels in d7 cells and the migratory and cranial neural crest markers *Sox9* and *Sox10* having the highest levels in d14 cells (Fig. 2A).

The strongest immunofluorescent staining was detected in d7 cells for Snai1 and d14 cells for Sox10 (Fig 2B).

**Fig 2.**
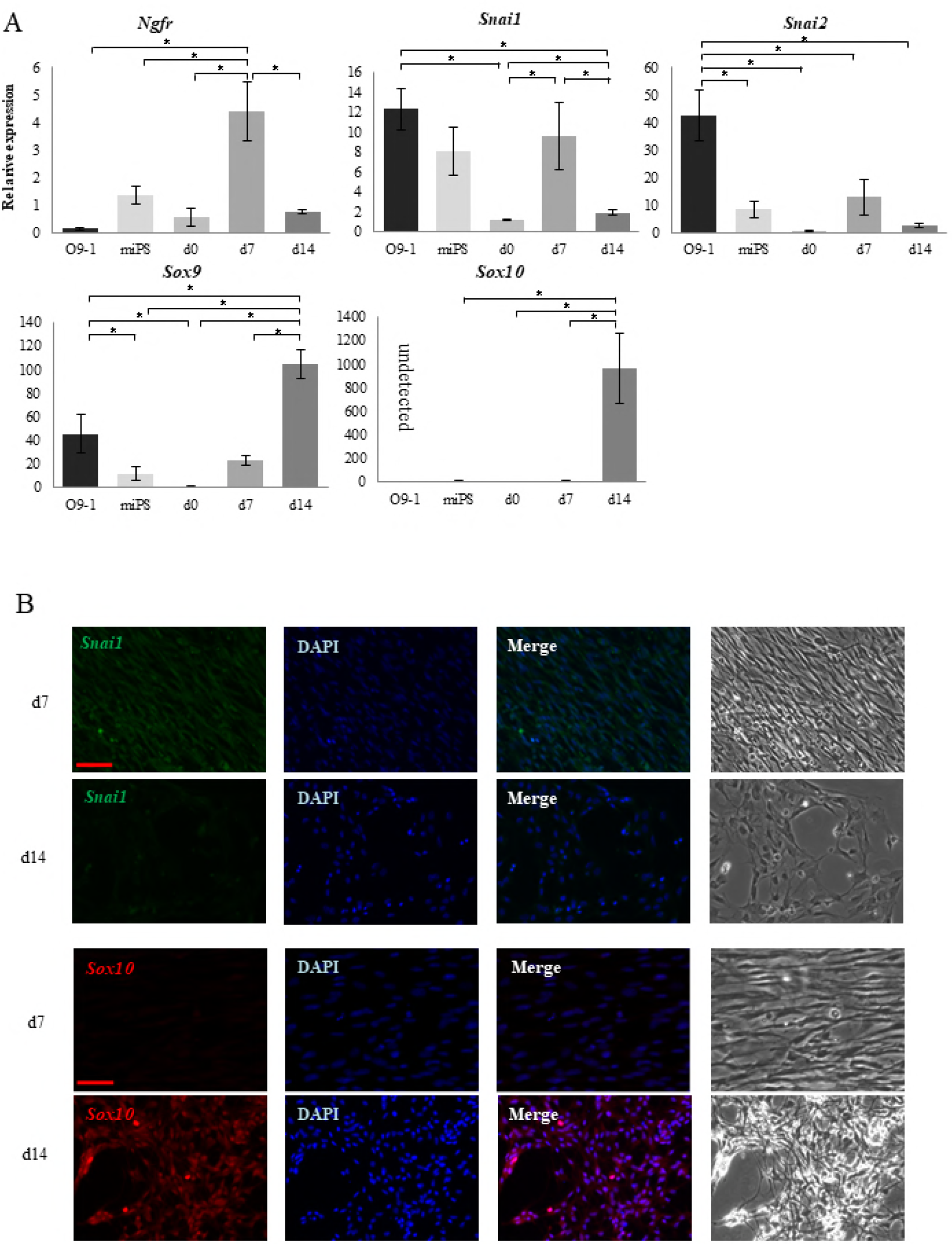
Comparison between O9–1 cells and cranial neural crest cells (cNCCs) derived from mouse induced pluripotent stem (miPS) cells using quantitative reverse transcription polymerase chain reaction (qRT-PCR) and immunostaining. (A) Expression of the premigratory neural crest (NC) markers *Ngfr, Snai1*, and *Snai2* and the migratory NC and cNC markers *Sox9* and *Sox10.* Expressions of the premigratory NC markers increased in day 7 (d7) cells, whereas those of the migratory markers increased in d14 cells. *Sox10* was not detected in the O9–1 cells. Each experiment was performed in triplicate with values representing the mean ± SD. Groups were compared using ANOVA followed by the Bonferroni test: *p < 0.05. (B) Immunostaining of d7 and d14 cells. Sox10 was more highly expressed in d14 cells, whereas Snai1

### NC specifier transcription factors

We conducted a literature search of NC specifier transcription factors that have been identified *in vivo* [16, 43–52, 60–106] (Tables 2 and 3) and compared these with our RNA-seq results. The relative expressions of genes that underwent a significant change in expression are presented in Fig 3A.

**Table 2.**
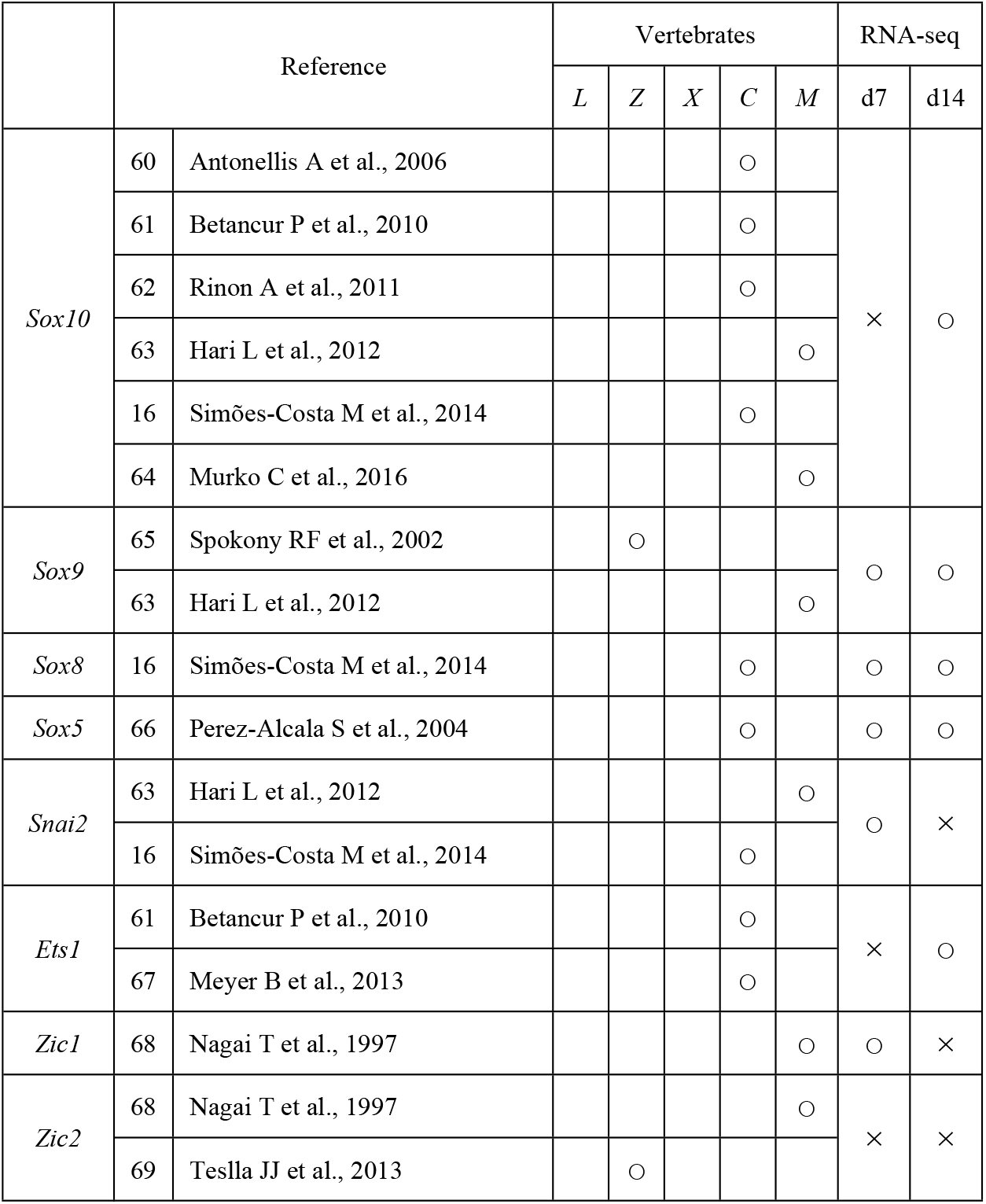
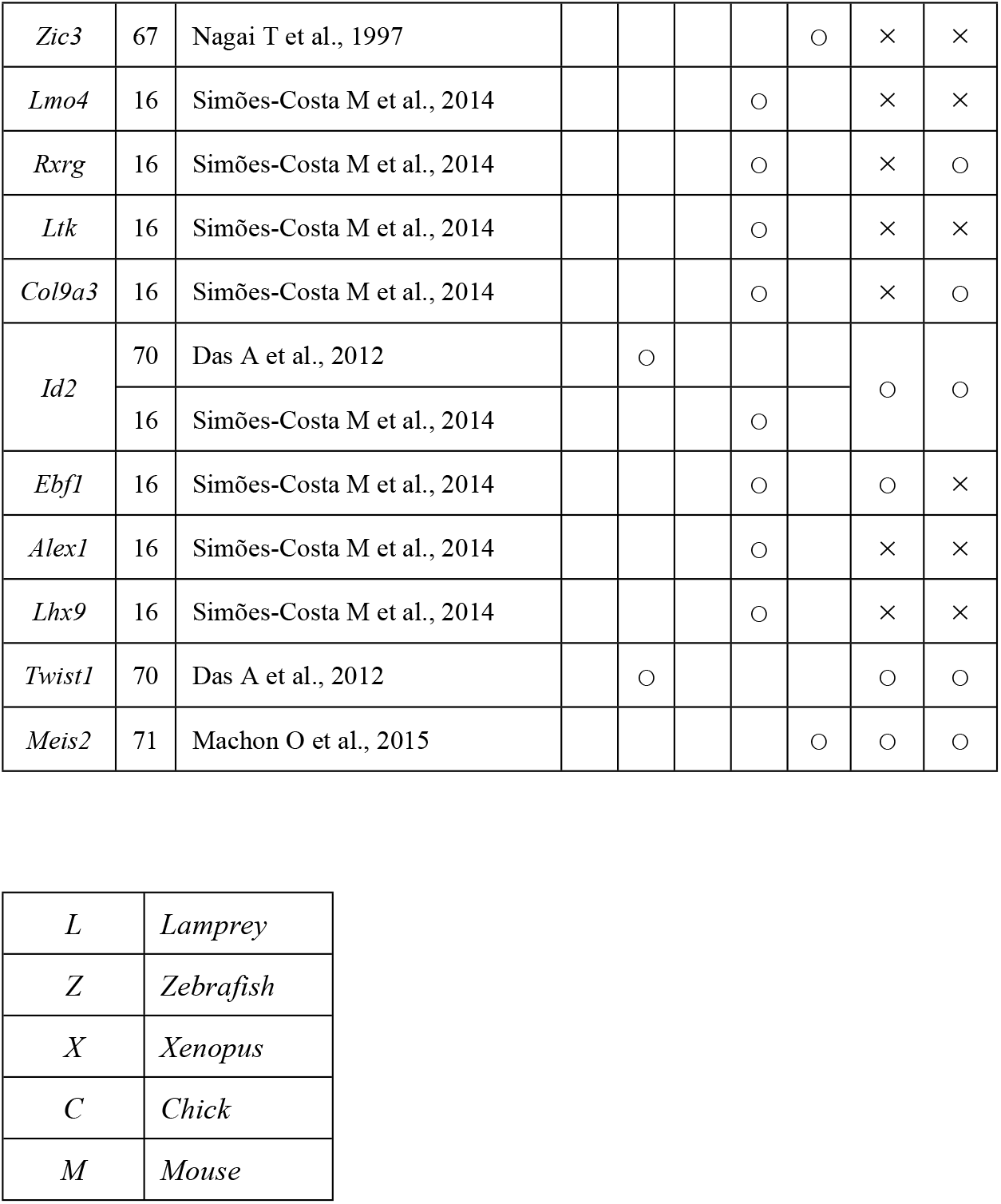
Cranial neural crest cells (cNCCs) genes that have previously been examined in vivo.

Open circles indicate genes that were upregulated on day 7 (d7) or d14 compared with d0 [log fold change (FC) > 1, *p* < 0.01, false discovery rate (FDR) < 0.05), whereas crosses indicate genes that were not upregulated.

**Table 3.**
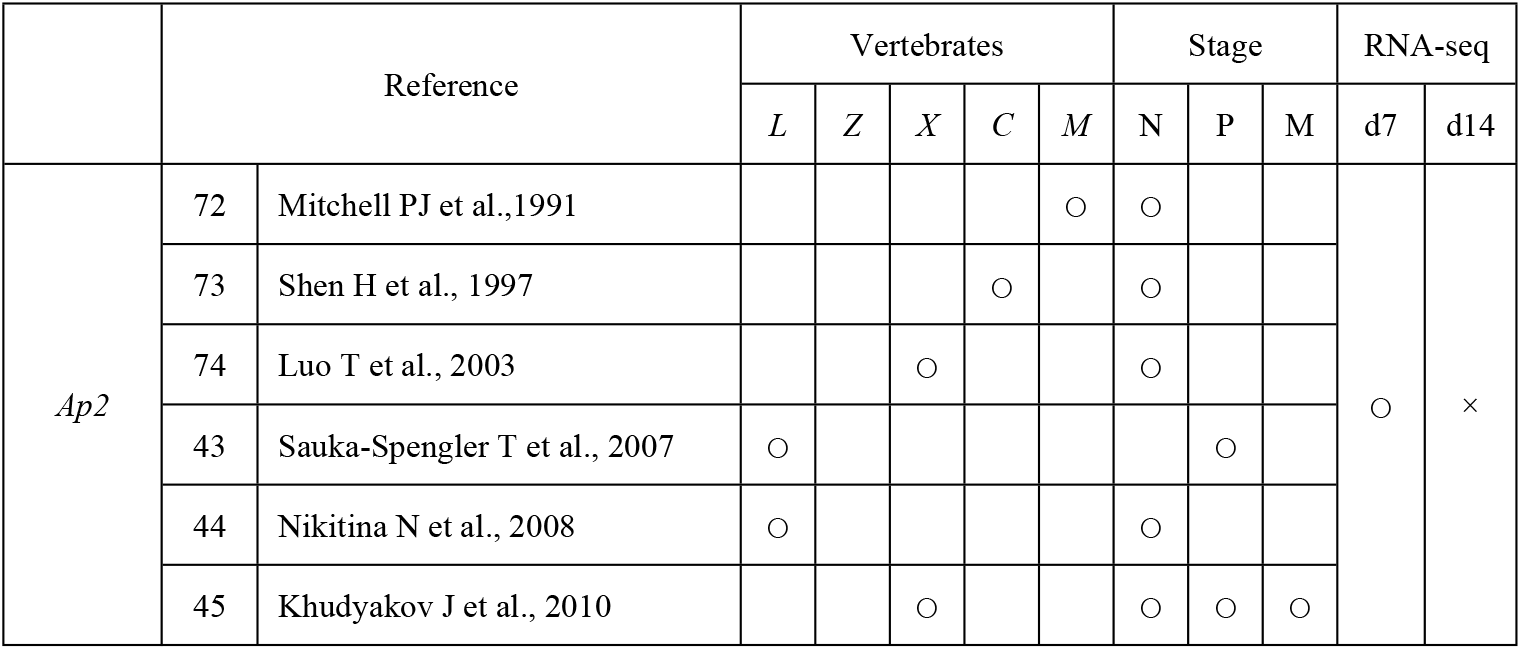
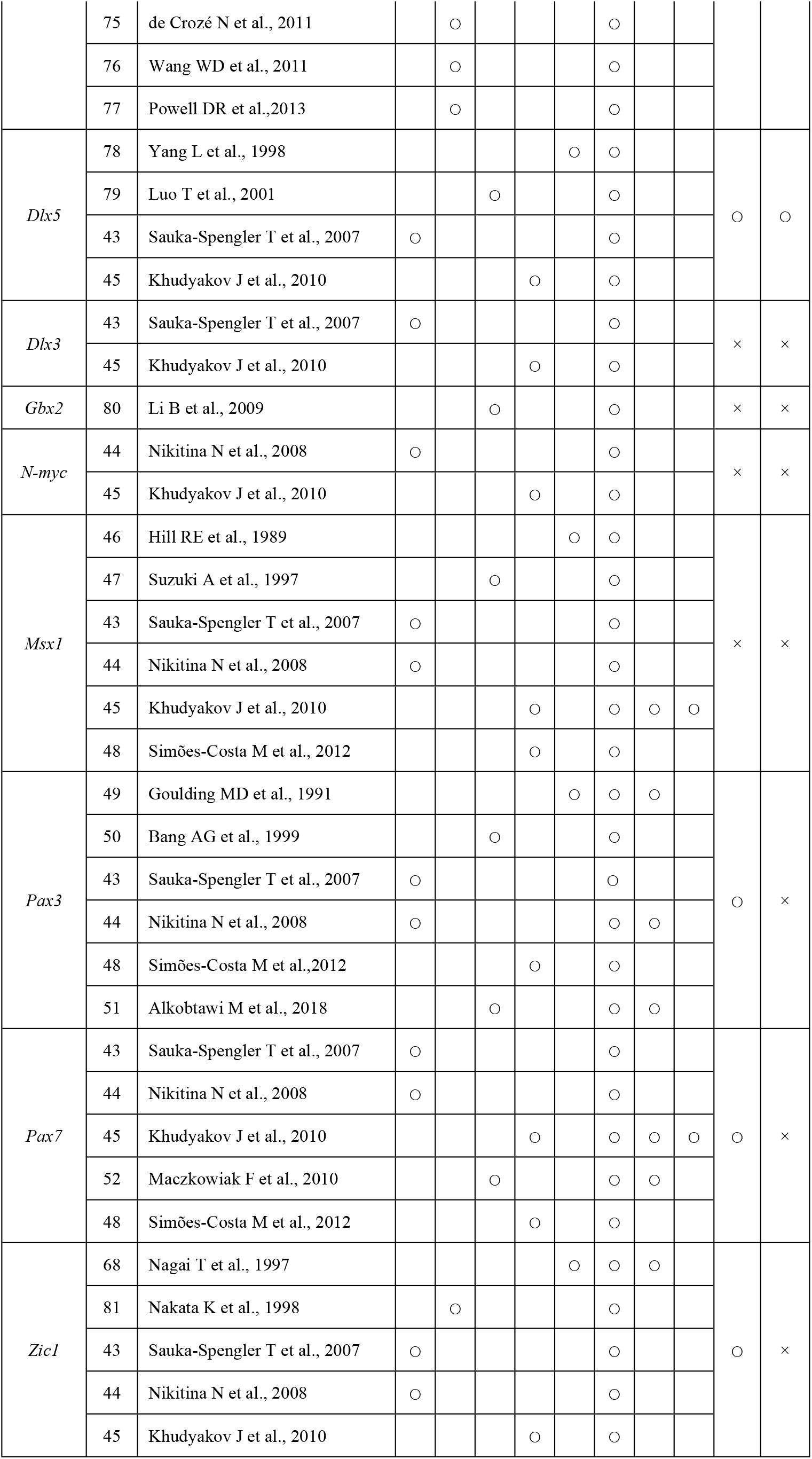
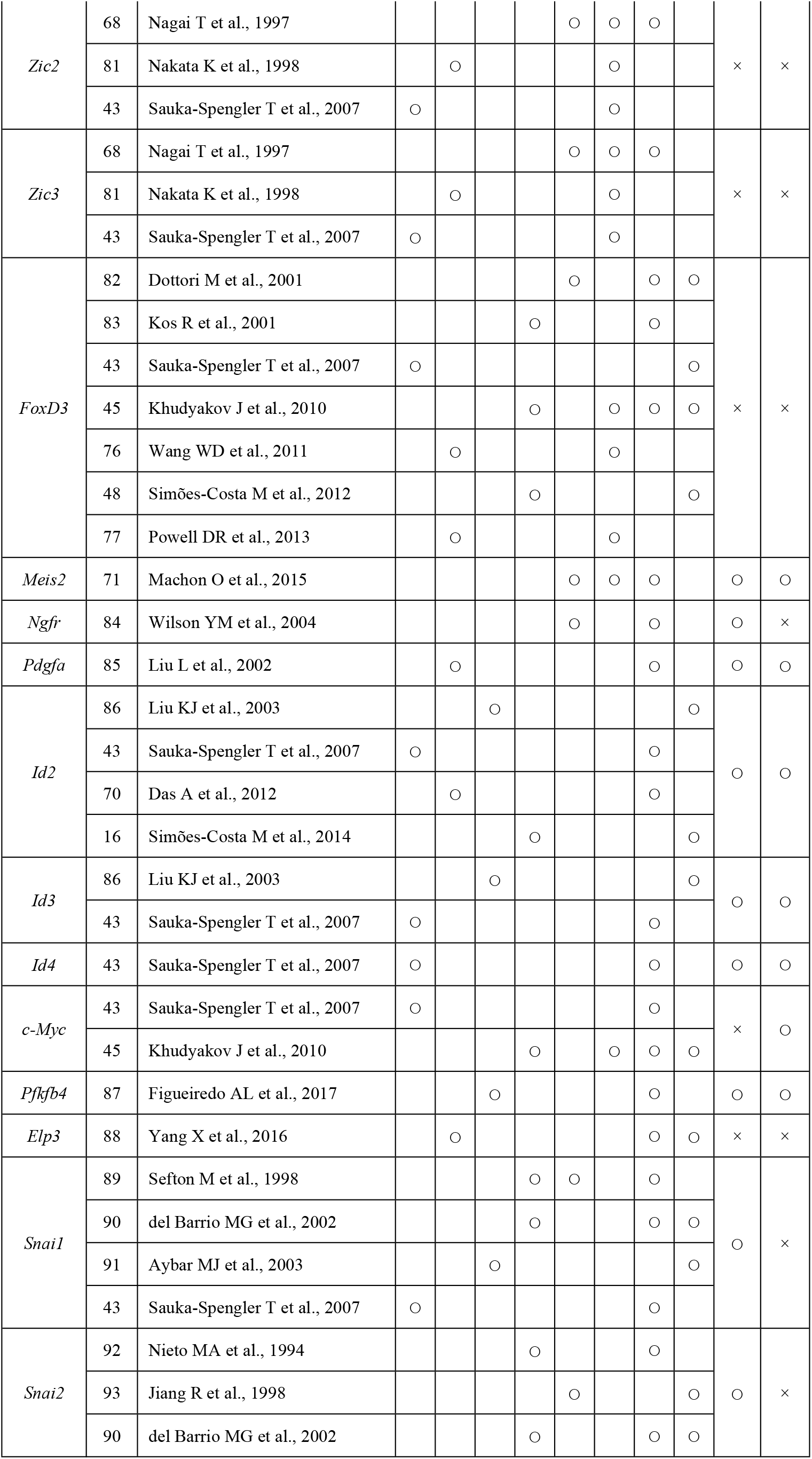
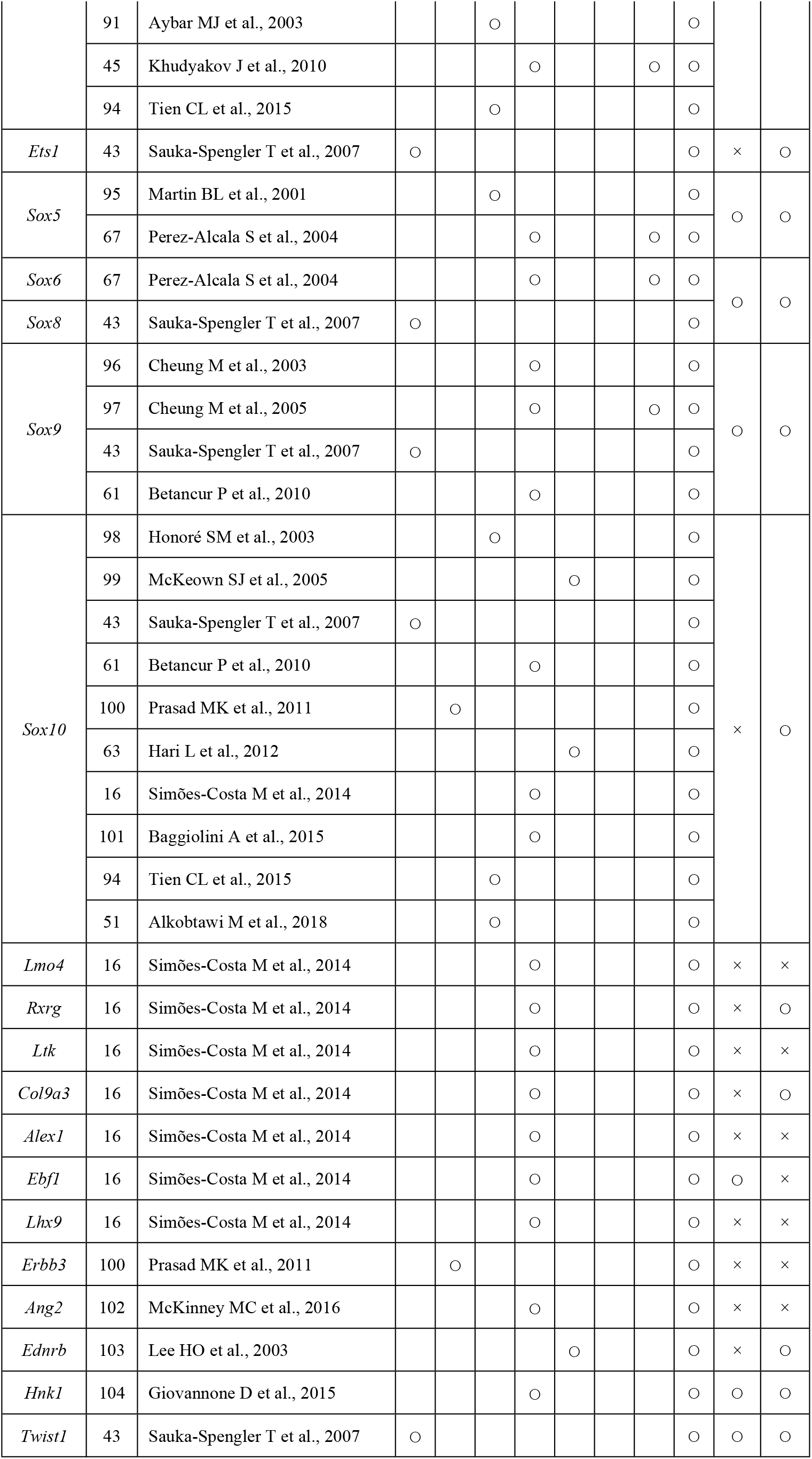
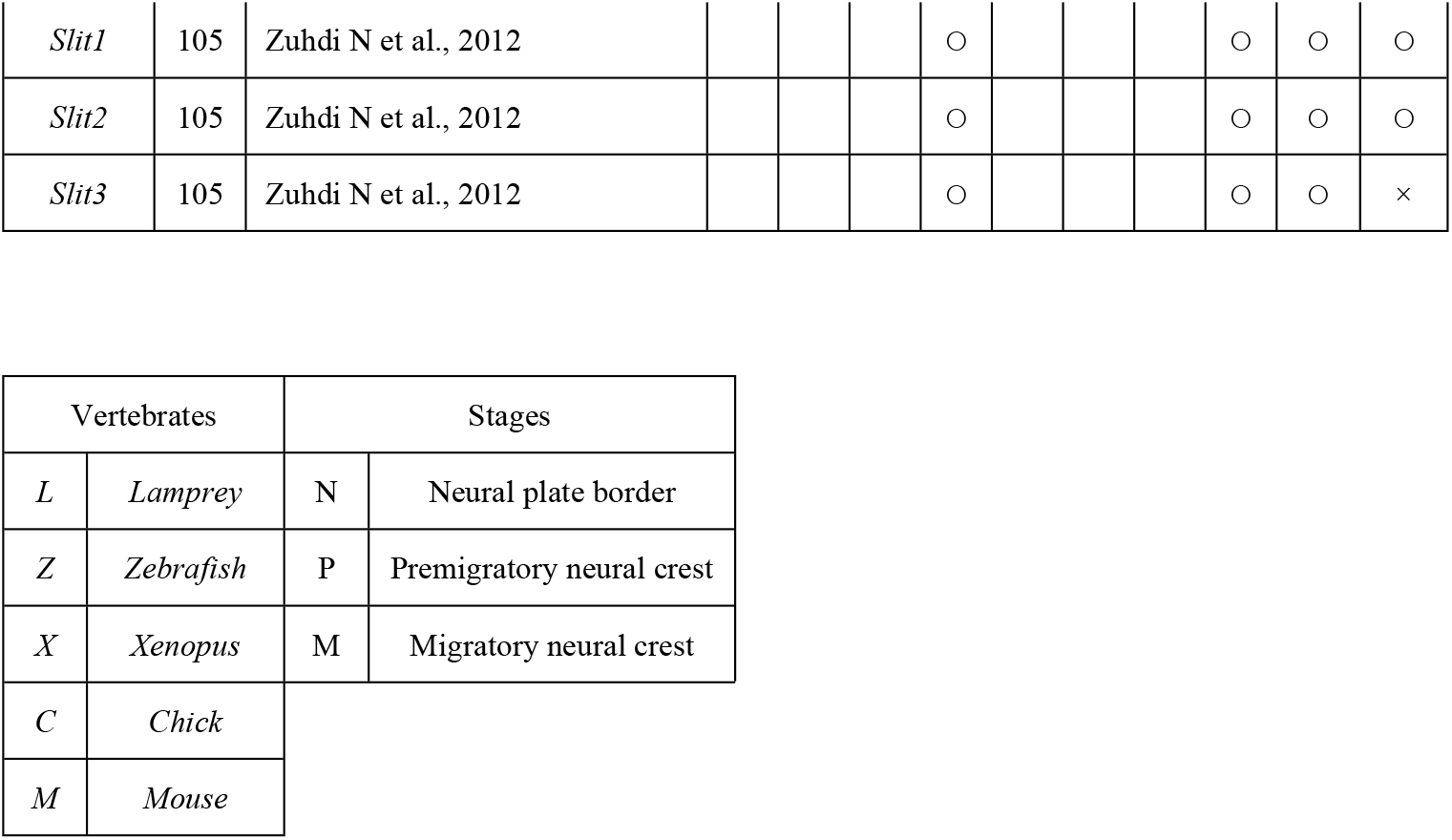
Neural crest (NC) transcription factors that have previously been examined *in vivo.*

Open circles indicate genes that were upregulated on day 7 (d7) or d14 compared with d0 [log fold change (FC) >1*, p* < 0.01, false discovery rate (FDR) < 0.05), whereas crosses indicate genes that were not upregulated.

**Fig 3.**
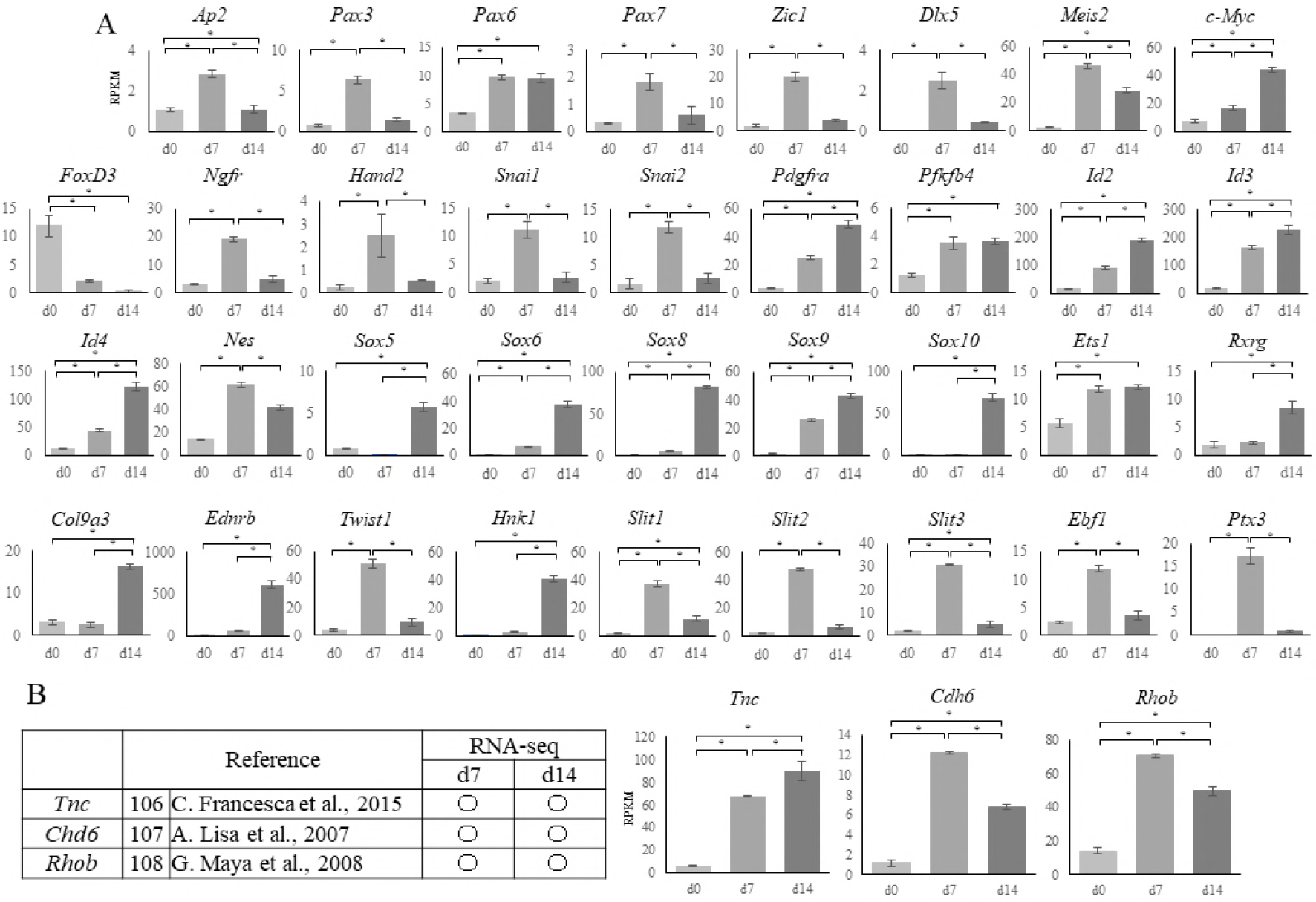
RNA sequencing results for cranial neural crest cells (cNCCs) differentiated from mouse induced pluripotent stem (miPS) cells. (A) Expression of each of the genes listed in Table 2 at day 0 (d0), d7, and d14 after induction. Sex-determining region Y (SRY)-related high mobility group (HMG) box genes were most upregulated in d14 cells. The vertical axis reveals reads per kilobase of exon per million mapped reads (RPKM) and the horizontal axis indicates time. Each experiment was performed in triplicate with values representing the mean ± SD. Groups were compared using ANOVA followed by the Bonferroni test: \**p* < 0.05. (B) Expression of genes that have not been examined during the neural crest stages *in vivo. Tnc* was most upregulated in the d14 cells, whereas *Cha6* and *Rhob* were upregulated in the d7 cells. The vertical axis reveals reads per kilobase of exon per million mapped reads (RPKM) and the horizontal axis indicates time. Open circles indicate genes that were upregulated on day 7 (d7) or d14 compared with d0 [log fold change (FC) > 1,*p* < 0.01, false discovery rate (FDR) < 0.05). Each experiment was performed in triplicate with values representing the mean ± SD. Groups were compared using ANOVA followed by the Bonferroni test: \**p* < 0.05.

We found that the *transcription factor AP-2 alpha* (*Ap2*) along with *Pax3* and zinc finger protein of the cerebellum 1 (*Zic1*), both of which are regulated by *Ap2*, were most highly expressed in d7 cells (Fig 3A). *Pax6*, which has been reported in human ES and iPS-derived NC cells (Tables 2 and 3), was detected in both d7 and d14 cells, whereas *Pax7*, which has not previously been reported in the mouse NC, was also detected in the d7 cells (Fig 3A). In contrast, the homeobox genes *gastrulation brain homeobox 2* (*Gbx2*)*, Msx1, distal-less homeobox 3* (*Dlx3*)*, Zic2*, and *Zic3* were not detected in d7 or d14 cells, and the homeobox genes *Zic1* and *Dlx5* were only expressed in d7 cells, despite these having been reported in the NC of a range of species (Table 2); however, *Meis homeobox 2* (*Meis2*) was expressed in both d7 and d14 cells.

Both MYCN proto-oncogene, bHLH transcription factor (*N-myc*) and *c-Myc* have been reported in NCCs (Table 3); however, we did not observe *N-myc* expression in d7 or d14 cells and detected *c-Myc* expression in the d7 and d14 group (Fig 3A). Furthermore, we observed substantial downregulation of the winged-helix transcription factor *FoxD3* over time (Fig 3A), which is an important factor for maintaining the pluripotency of ES cells and a key NC specifier that has been implicated in multiple steps of NC development and NCC migration in the embryo of various species (Table 2).

The premigratory NC markers *Ngfr*, heart and neural crest derivatives expressed 2 (*Hand2*)*, Snai1*, and *Snai2* were only detected in the d7 cells; however, other premigratory NC markers, such as platelet derived growth factor receptor, alpha polypeptide (*Pdgfra*), 6-phosphofructo-2-kinase/fructose-2,6-biphosphatase 4 (*Pfkfb4*), inhibitor of DNA binding 2 (*Id2*)*, Id3*, and *Id4* were found in both d7 and d14 cells, as was *Nes* (Fig 3A).

Migratory neural crest markers expression of *Sox5, -6, -8, -9*, and *-10*, which encode members of the sex-determining region Y (SRY)-related high mobility group (HMG)-box family of transcription factors and have been reported to be crucial in several aspects of NCCs, were observed in d7 or d14 cells. *Sox10*, which is a known marker of migratory cNCCs in various species (Table 2), was only detected in d14 cells, as were other migratory NC markers, including retinoid X receptor gamma (*Rxrg*), collagen type IX alpha 3 chain (*Col9a3*), and endothelin receptor type B (*Ednrb;* Fig. 3A); however, LIM domain only 4 (*Lmo4*), leukocyte tyrosine kinase (*Ltk*), erb-b2 receptor tyrosine kinase 3 (*Erbb3*), and angiogenin, ribonuclease A family, member 2 (*Ang2*) were not detected in the d14 cells.

Twist family bHLH transcription factor 1 (*Twist1*), which is activated by a variety of signal transduction pathways and is crucial in the downregulation of E-cadherin expression, was detected in both d7 and d14 cells, as was beta-1,3-glucuronyltransferase 1 (*B3gat1/Hnk1*), or CD57. In contrast, expression of the trunk NC markers lit guidance ligand 1/2 (*Slit1/2*), which has been reported to play an important role in the migration of trunk NC cells toward ventral sites, was upregulated only in the d7 cells (Fig 3A).

Eventually, the expressions of tenascin C (Tnc), cadherin-6 (*Cdh6*), and ras homolog family member B (*Rhob*), all of which are related to cell adhesion and motility [106–111], significantly increased in both d7 and d14 cells (Fig 3B).

### Metzincin superfamily zinc proteinase and protocadherin superfamily expressions

Members of the metzincin superfamily are proteinases that have a zinc ion at their active site. This family includes the matrix metalloproteinases (Mmps), a disintegrin and metalloproteinase (Adam), and Adamts, all of which have attracted attention as factors involved in cancer cell invasion and cell migration. *Mmp2, -11, -14, -15, -16, -24*, and *-28* were significantly upregulated in the cNNCs (Fig 4A), all of which except *Mmp24* are membrane-bound types. The expressions of *Mmp11* and *-28* were only detected in d7 cells, while all other Mmps were detected in both d7 and d14 cells (Fig 4A, B).

Only *Adam1a, -8, -10*, and *-12* were upregulated in both d7 and d14 cells (Fig 4C, D), despite the members of this family being important in NC migration and the expressions of *Adam10, -12, -15, -19*, and-33 having been observed in the mouse NC [112]. In contrast, various *Adamts* family genes, which are important for connective tissue organization and cell migration, were upregulated in either d7 or d14 cells (Fig 4C, D). The expression of *Adamts1* in particular exhibited a substantial increase in expression, while *Adamts2* and *-8*, which are presumed to be important in cancer cell invasion [55], increased in the later stages of differentiation.

**Fig. 4.**
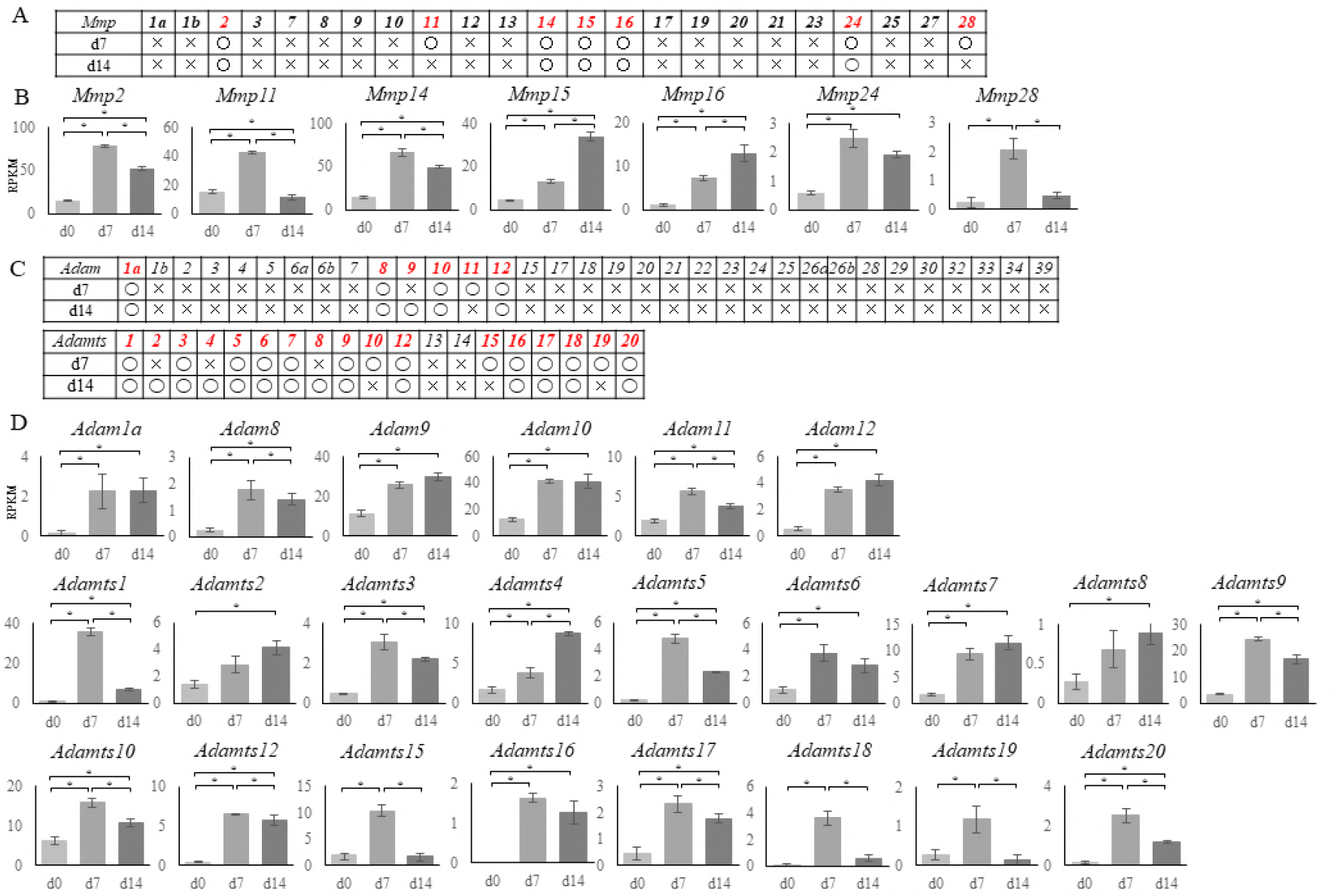
RNA sequencing results for the matrix metalloproteinase (*Mmp*), a disintegrin and metalloproteinase (*Adam*), and a disintegrin and metalloproteinase with thrombospondin motifs (*Adamts*) gene families. (A) Expressions of *Mmp* family genes in mouse. Round marks alongside d7 or d14 indicate that the genes were upregulated compared with d0 [log fold change (logFC) > 1, *p* < 0.01, false discovery rate (FDR) <0.05], whereas cross marks indicate no upregulation in d7 or d14 cells. (B) Graphical representation of the upregulation of *Mmp2, -11, - 14, -15, -16, -24*, and *-28* in d7 or d14 cells. *Mmp15* and *-16* were most upregulated in d14 cells.

The vertical axis reveals reads per kilobase of exon per million mapped reads (RPKM) and the horizontal axis indicates time. Each experiment was performed in triplicate with values representing the mean ± SD. Groups were compared using ANOVA followed by the Bonferroni test: \**p* < 0.05. (C) Expressions of *Adam* and *Adamts* genes in mouse. Round marks alongside d7 or d14 indicate that the genes were upregulated compared with d0 (logFC > 1, p < 0.01, FDR < 0.05), whereas cross marks indicate no upregulation. (D) Graphical representation of the upregulation of *Adam1a* and *8–12*, and *Adamts1–10, -12*, and *15–20* in d7 or d14 cells. *Adam2, -4, -7*, and *-8*, and *Adamts 9* and *-12* were most upregulated in d14 cells. The vertical axis reveals reads per kilobase of exon per million mapped reads (RPKM) and the horizontal axis indicates time. Each experiment was performed in triplicate with values representing the mean ± SD. Groups were compared using ANOVA followed by the Bonferroni test: \**p* < 0.05.

Most of the *Pcdh* genes, which are involved in cell adhesion, were upregulated in d7 and d14 cells (Table 4); however, *Pcdha2, -5, -7, -11*, and *-12; Pcdhac1;* and *Pcdhgc5* were only upregulated in the d14 cells.

**Table 4.**
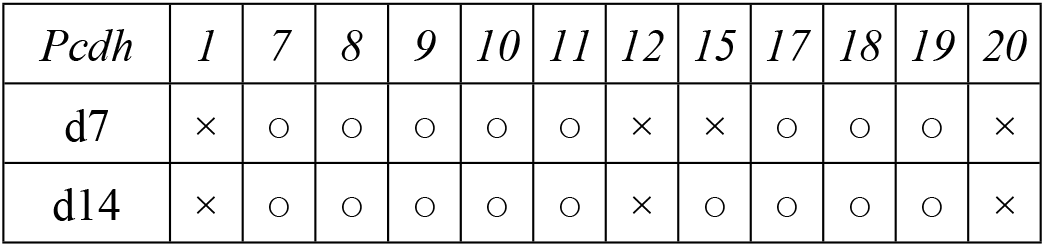
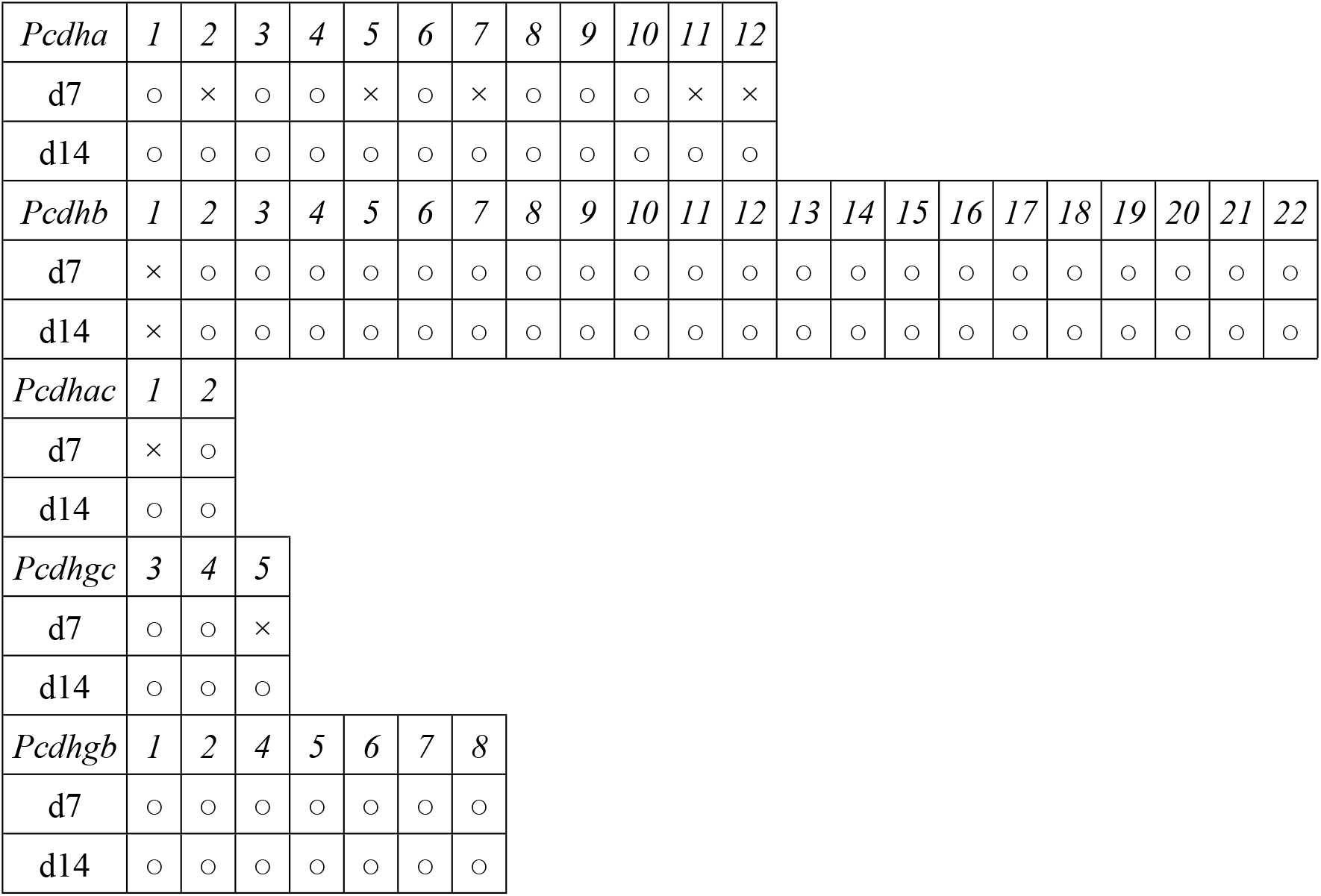
Expression of the protocadherin superfamily based on RNA sequencing data. Open circles indicate genes that were upregulated on day 7 (d7) or d14 compared with d0 [log fold change (FC) >1*, p* < 0.01, false discovery rate (FDR) < 0.05), whereas crosses indicate genes that were not upregulated.

| *Pcdh* | 1 | 7 | 8 | 9 | 10 | 11 | 12 | 15 | 17 | 18 | 19 | 20 |
| --- | --- | --- | --- | --- | --- | --- | --- | --- | --- | --- | --- | --- |
| d7 | × | ○ | ○ | ○ | ○ | ○ | × | × | ○ | ○ | ○ | × |
| d14 | × | ○ | ○ | ○ | ○ | ○ | × | ○ | ○ | ○ | ○ | × |

|  |  |  |  |  |  |  |  |  |  |  |  |  |  |  |  |  |  |  |  |  |  |  |
| --- | --- | --- | --- | --- | --- | --- | --- | --- | --- | --- | --- | --- | --- | --- | --- | --- | --- | --- | --- | --- | --- | --- |
| *Pcdha* | 1 | 2 | 3 | 4 | 5 | 6 | 7 | 8 | 9 | 10 | 11 | 12 |  |  |  |  |  |  |  |  |  |  |
| d7 | ○ | × | ○ | ○ | × | ○ | × | ○ | ○ | ○ | × | × |  |  |  |  |  |  |  |  |  |  |
| d14 | ○ | ○ | ○ | ○ | ○ | ○ | ○ | ○ | ○ | ○ | ○ | ○ |  |  |  |  |  |  |  |  |  |  |
| *Pcdhb* | 1 | 2 | 3 | 4 | 5 | 6 | 7 | 8 | 9 | 10 | 11 | 12 | 13 | 14 | 15 | 16 | 17 | 18 | 19 | 20 | 21 | 22 |
| d7 | × | ○ | ○ | ○ | ○ | ○ | ○ | ○ | ○ | ○ | ○ | ○ | ○ | ○ | ○ | ○ | ○ | ○ | ○ | ○ | ○ | ○ |
| d14 | × | ○ | ○ | ○ | ○ | ○ | ○ | ○ | ○ | ○ | ○ | ○ | ○ | ○ | ○ | ○ | ○ | ○ | ○ | ○ | ○ | ○ |
| *Pcdhac* | 1 | 2 |  |  |  |  |  |  |  |  |  |  |  |  |  |  |  |  |  |  |  |  |
| d7 | × | ○ |  |  |  |  |  |  |  |  |  |  |  |  |  |  |  |  |  |  |  |  |
| d14 | ○ | ○ |  |  |  |  |  |  |  |  |  |  |  |  |  |  |  |  |  |  |  |  |
| *Pcdhgc* | 3 | 4 | 5 |  |  |  |  |  |  |  |  |  |  |  |  |  |  |  |  |  |  |  |
| d7 | ○ | ○ | × |  |  |  |  |  |  |  |  |  |  |  |  |  |  |  |  |  |  |  |
| d14 | ○ | ○ | ○ |  |  |  |  |  |  |  |  |  |  |  |  |  |  |  |  |  |  |  |
| *Pcdhgb* | 1 | 2 | 4 | 5 | 6 | 7 | 8 |  |  |  |  |  |  |  |  |  |  |  |  |  |  |  |
| d7 | ○ | ○ | ○ | ○ | ○ | ○ | ○ |  |  |  |  |  |  |  |  |  |  |  |  |  |  |  |
| d14 | ○ | ○ | ○ | ○ | ○ | ○ | ○ |  |  |  |  |  |  |  |  |  |  |  |  |  |  |  |

## Discussion

In this study, we successfully generated miPS-induced cNCCs that were sufficiently close to the migratory stage. The NC has previously been generated from ES or iPS cells in various ways [24,33–39] and the protocol we used in the present study was based on the methods outlined by R. Bajpai et al. [39]; however, few studies have investigated the changes in the properties of cNCCs at different time points (Table 5).

**Table 5.**
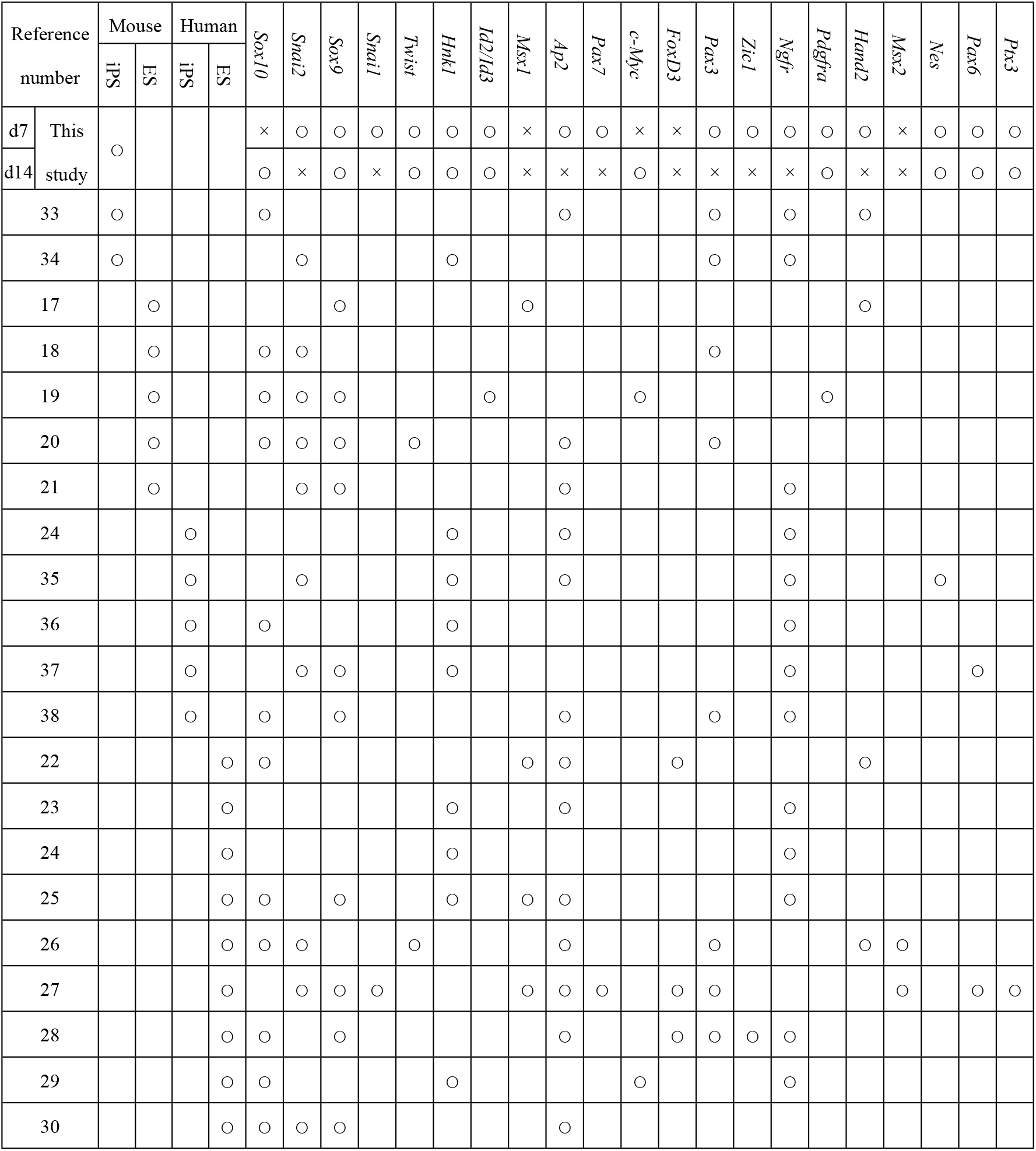
Neural crest (NC) transcription factors that have previously been examined *in vitro.* Open circles indicate genes that were upregulated on day 7 (d7) or d14 compared with d0 [log fold change (FC) >1*, p* < 0.01, false discovery rate (FDR) < 0.05), whereas crosses indicate genes that wer not upregulated.

Our d7 and d14 cells expressed typical NC markers, such as *Ngfr, Snai1*, and *Snai2.* In contrast, the mouse cNCC line (O9–1 cells) did not express *Ngfr*, indicating that cNCCs derived from miPS cells may be of better quality for evaluating the cNCC characteristics than O9–1 cells [59]. We also found that, unlike O9–1 cells, d14 cells expressed considerably high levels of *Sox10*, which is considered as a reliable marker for migratory cNCCs. Because cNCCs are involved in organizing numerous craniofacial tissues, several reports are available on their gene expression profiles; however, we found that various results that have been reported were inconsistent between the species and protocols. Since cNCCs differentiate fast in the embryo [14], it is considerably difficult to synchronize the timing of isolation to a particular point in their development. Furthermore, migratory cNCCs intermingle with other types of cells in the embryo, making it difficult to isolate and characterize a pure cell population. Consequently, there have been few reports of cNCC markers [16,60–71]; however, Simoes-Costa et al. [16] successfully isolated *Sox10* positive cNCCs in a chicken embryo and analyzed their gene profiles and we found that d14 cells expressed several of these *Sox10* positive chicken cNCCs. It has previously been suggested that NC cells have multiple populations [11] and that the generation of cNCCs from iPS cells could result in numerous different populations occurring in the same dish. Therefore, this diversity in populations may explain the discrepancies; however, we can conclude that under the conditions used in the present study, *cMyc; Ets1; Sox10; Adamts2; Adamts8; Pcdha2, -5, -7, -*11, and-12; *Pcdhac1*, and *Pcdhgc3* may represent useful markers for migratory cNCCs.

Our results also indicated that d7 cells were still in the premigratory stage even though they expressed numerous NC markers. Thus, cNCCs derived from miPS cells took more than 14 days to become migratory *in vitro*, which is much slower than has been observed in the mouse embryos *in vivo* under the same conditions [113].

RNA-seq makes it possible to normalize the expression levels of different genes, allowing comparisons between samples. We conducted triplicate experiments in which none of the induced cNCCs expressed several homeobox genes that are considered to be expressed in the early stages of cNCC differentiation. In particular, we did not observe *FoxD3* expression in either d7 or d14 cells, despite it being recognized as one of the key transcription factors in cNCCs [53]. These negative results indicate that cNCCs derived from miPS cells may have distinct gene regulatory networks. Although it is possible that the cells would express those genes at different time points, the expression of *FoxD3*, which is a pluripotent stem cell marker gene and plays an important role in maintaining pluripotency, decreases in a time-dependent manner [44], making it more likely that *FoxD3* may not be a key regulator in iPS-derived cNCCs. We speculate, however, that iPS cells had a sufficient amount of *FoxD3* to allow them to be converted from iPS cells into cNCCs.

Protocadherins belong to the cadherin superfamily and are involved in intercellular interactions [57], while metzincins are thought to be key proteinases that facilitate the cell migration [45]. Unfortunately, the abundances of members of these families hindered their analysis; however, since RNA-seq techniques enable us to evaluate the gene profiles exhaustively, we were able to focus on the expressions of all of the procadherin and metazicin family members. As expected, we found that several *Adam* and *Adamts* genes were upregulated, with most of the latter increasing significantly. The *Adam* genes that increased in the cNCCs were the membrane-bound type; whereas, the *Adamts* genes were secreted proteinases, indicating that the expression of various *Adamts* may allow the matrix to be digested more efficiently, as each may be capable of digesting a different type of extracellular matrix protein [45]. Thus, the secretion of a variety of Adamts and Pcdh proteins may play a crucial role in the migration ability of cNCCs.

In summary, we successfully induced the formation of cNCCs from miPS cells by placing them in NC inducing media for 14 days. We found that although the resulting cNCCs had several NC specifiers, some were lacking, indicating that a distinct molecular network may control the gene expression in miPS-derived cNCCs. Our results also indicated that *cMyc; Ets1; Sox10; Adamts2* and *-8; Pcdha2, -5, -7*, *-11*, and *-12; Pcdhac1;* and *Pcdhgc3* may represent appropriate markers for migratory cNCCs induced from miPS cells. Eventually, these cNCCs produced a broad spectrum of Adamts family proteins that may play an important role in their migration.

## Acknowledgments

The author is grateful to Professor T. Azuma, MD, PhD, Department of Biochemistry, and Professor T. Ichinohe, DDS, PhD, Department of Dental Anesthesiology, for their guidance. I also thank S. Onodera and A. Saito, Department of Biochemistry.

## References

1. Luan X, Dangaria S, Ito Y, Walker CG, Jin T, Schmidt MK et al. Neural crest lineage segregation: a blueprint for periodontal regeneration. J Dent Res. 2009; 88: 781–791. https://doi.org/10.1177/0022034509340641 PMID: 19767574

2. Malhotra N. Induced Pluripotent Stem (iPS) Cells in Dentistry: A Review, Int J Stem Cells. 2016; 9: 176–185. https://doi.org/10.15283/ijsc16029 PMID: 27572712

3. Knight RD, Schilling TF. Cranial neural crest and development of the head skeleton. Adv Exp Med Biol. 2006; 589: 120–133. https://doi.org/10.1007/978-0-387-46954-67 PMID: 17076278

4. Theveneau E, Mayor R. Collective cell migration of the cephalic neural crest: the art of integrating information. Genesis. 2011; 49: 164–176. https://doi.org/10.1002/dvg.20700 PMID: 21157935

5. Chai Y, Jiang X, Ito Y, Bringas P Jr, Han J, Rowitch DH et al. Fate of the mammalian cranial neural crest during tooth and mandibular morphogenesis. Development.2000; 127: 1671–1679. PMID: 10725243

6. McConnell AM, Mito IK, Ablain J, Dang M, Formichella L, Fisher DE et al. Neural crest state activation in NRAS driven melanoma, but not in NRAS-driven melanocyte expansion. Dev Biol. 2018. https://doi.org/10.1016/j.ydbio.2018.05.026 PMID: 29883661

7. Meulemans D, Bronner-Fraser M. Gene-regulatory interactions in neural crest evolution and Development. Dev Cell. 2004; 7: 291–299. https://doi.org/10.1016/j.devcel.2004.08.007 PMID: 15363405

8. Steventon B, Carmona-Fontaine C, Mayor R. Genetic network during neural crest induction: from cell specification to cell survival. Semin Cell Dev Biol. 2005; 16: 647–654. https://doi.org/10.1016/j.semcdb.2005.06.001 PMID: 16084743

9. Simões-Costa M, Bronner ME. Establishing neural crest identity: a gene regulatory recipe. Development. 2015; 142: 242–257. https://dx.doi.org/10.1242%2Fdev.105445 PMID: 25564621

10. Martik ML, Bronner ME. Regulatory Logic Underlying Diversification of the Neural Crest. Trends Genet. 2017; 33: 715–727. https://doi.org/10.1016/j.tig.2017.07.015 PMID: 28851604

11. Minoux M, Rijli FM. Molecular mechanisms of cranial neural crest cell migration and patterning in craniofacial development. Development. 2010; 137: 2605–2621. https://doi.org/10.1242/dev.040048 PMID: 20663816

12. Mayor R, Theveneau E. The neural crest. Development. 2013; 140: 2247–2251. https://doi.org/10.1242/dev.091751 PMID: 23674598

13. Okuno H, Mihara FR, Ohta S, Fukuda K, Kurosawa K, Akamatsu W et al. CHARGE syndrome modeling using patient-iPSCs reveals defective migration of neural crest cells harboring CHD7 mutations. eLife. 2017; 6: e21114 https://dx.doi.org/10.7554%2FeLife.21114 PMID: 29179815

14. Simoes-Costa M, Bronner ME. Reprogramming of avian neural crest axial identity and cell fate. Science. 2016; 352: 1570–1573. https://dx.doi.org/10.1126%2Fscience.aaf2729 PMID: 27339986

15. Milet C, Monsoro Burq AH. Neural crest induction at the neural plate border in vertebrates. Dev Biol. 2012; 366: 22–33. https://doi.org/10.1016/j.ydbio.2012.01.013 PMID: 22305800

16. Simões-Costa M, Tan-Cabugao J, Antoshechkin I, Sauka-Spengler T, Bronner ME. Transcriptome analysis reveals novel players in the cranial neural crest gene regulatory Network. Genome Res. 2014; 24: 281–290. https://doi.org/10.1101/gr.161182.113 PMID: 24389048

17. Mizuseki K, Sakamoto T, Watanabe K, Muguruma K, Ikeya M, Nishiyama A et al. Generation of neural crest-derived peripheral neurons and floor plate cells from mouse and primate embryonic stem cells. Proc Natl Acad Sci U S A. 2003; 100: 5828–5833. https://dx.doi.org/10.1073%2Fpnas.1037282100 PMID: 12724518

18. Motohashi T, Aoki H, Chiba K, Yoshimura N, Kunisada T. Multipotent Cell Fate of Neural Crest-Like Cells Derived from Embryonic Stem Cells. Stem Cells. 2007; 25: 402–412. https://doi.org/10.1634/stemcells.2006-0323 PMID: 17038669

19. Kawaguchi J, Nichols J, Gierl MS, Faial T, Smith A. Isolation and propagation of enteric neural crest progenitor cells from mouse embryonic stem cells and embryos. Development. 2010; 137: 693–704. https://dx.doi.org/10.1242%2Fdev.046896 PMID: 20147374

20. Aihara Y, Hayashi Y, Hirata M, Ariki N, Shibata S, Nagoshi N et al. Furue, Induction of neural crest cells from mouse embryonic stem cells in a serum-free monolayer culture. Int J Dev Biol. 2010; 154: 1287–1294. https://doi.org/10.1387/ijdb.103173ya PMID: 20711997

21. Minamino Y, Ohnishi Y, Kakudo K, Nozaki M. Isolation and propagation of neural crest stem cells from mouse embryonic stem cells via cranial neurospheres. Stem Cells Dev. 2015; 24: 172–181. https://doi.org/10.1089/scd.2014.0152 PMID: 25141025

22. Pomp O, Brokhman I, Ben-Dor I, Reubinoff B, Goldstein RS. Generation of peripheral sensory and sympathetic neurons and neural crest cells from human embryonic stem cells. Stem cells. 2005; 23: 923–930. https://doi.org/10.1634/stemcells.2005-0038 PMID: 15883233

23. Lee G, Kim H, Elkabetz Y, Al Shamy G, Panagiotakos G, Barberi T et al. Isolation and directed differentiation of neural crest stem cells derived from human embryonic stem cells. Nat Biotechnol. 2007; 25: 1468–1475. https://doi.org/10.1038/nbt1365 PMID: 18037878

24. Lee G, Chambers SM, Tomishima MJ, Studer. Derivation of neural crest cells from human pluripotent stem cells. Nat Protoc. 5 (2010) 688–701. https://doi.org/10.1038/nprot.2010.35 PMID: 20360764

25. Liu Q, Spusta SC, Mi R, Lassiter RN, Stark MR, Höke A et al. Human neural crest stem cells derived from human ESCs and induced pluripotent stem cells: induction, maintenance, and differentiation into functional schwann cells. Stem Cells Transl Med. 2010; 1: 266–278. https://doi.org/10.5966/sctm.2011-0042 PMID: 23197806

26. Noisa P, Lund C, Kanduri K, Lund R, Lähdesmäki H, Lahesmaa R et al. Notch signaling regulates the differentiation of neural crest from human pluripotent stem cells. J Cell Sci. 2014; 127: 2083–2094. https://doi.org/10.1242/jcs.145755 PMID: 24569875

27. Karbalaie K, Tanhaei S, Rabiei F, Kiani-Esfahani A, Masoudi NS, Nasr-Esfahani MH et al. Stem cells from human exfoliated deciduous tooth exhibit stromal-derived inducing activity and lead to generation of neural crest cells from human embryonic stem cells. Cell J. 2015; 17: 37–48. https://dx.doi.org/10.22074%2Fcellj.2015.510 PMID: 25870833

28. Avery J, Dalton S. Methods for Derivation of Multipotent Neural Crest Cells Derived from Human Pluripotent Stem Cells. Methods Mol Biol. 2016; 1341: 197–208. https://dx.doi.org/10.1007%2F7651_2015_234 PMID: 25986498

29. Zhang JT, Weng ZH, Tsang KS, Tsang LL, Chan HC, Jiang XH. MycN Is Critical for the Maintenance of Human Embryonic Stem Cell-Derived Neural Crest Stem Cells. PLoS One; 2016: e0148062. https://doi.org/10.1371/journal.pone.0148062 PMID: 26815535

30. Lovatt M, Yam GH, Peh GS, Colman A, Dunn NR, Mehta JS. Directed differentiation of periocular mesenchyme from human embryonic stem cells. Differentiation. 2018; 99: 62–69. https://doi.org/10.1016/j.diff.2017.11.003 PMID: 29239730

31. Doi D, Samata B, Katsukawa M, Kikuchi T, Morizane A, Ono Y et al. Isolation of human induced pluripotent stem cell derived dopaminergic progenitors by cell sorting for successful transplantation. Stem Cell Reports. 2014; 3: 337–350. https://dx.doi.org/10.1016%2Fj.stemcr.2014.01.013 PMID: 24672756

32. Nakane T, Masumoto H, Tinney JP, Yuan F, Kowalski WJ, Ye F et al. Impact of Cell Composition and Geometry on Human Induced Pluripotent Stem Cells-Derived Engineered Cardiac Tissue. Sci Rep. 2017; 7: 45641. https://dx.doi.org/10.1038%2Fsrep45641 PMID: 28368043

33. Okawa T, Kamiya H, Himeno T, Kato J, Seino Y, Fujiya A. Transplantation of Neural Crest Like Cells Derived From Induced Pluripotent Stem Cells Improves Diabetic Polyneuropathy in Mice. Cell Transplant. 2013; 22: 1767–1783. https://doi.org/10.3727/096368912X657710 PMID: 23051637

34. Seki D, Takeshita N, Oyanagi T, Sasaki S, Takano I, Hasegawa M et al. Differentiation of Odontoblast-Like Cells From Mouse Induced Pluripotent Stem Cells by Pax9 and Bmp4 Transfection Stem Cells. Transl Med. 2015: 4: 993–997. https://dx.doi.org/10.5966%2Fsctm.2014-0292 PMID: 26136503

35. Wang A, Tang X, Li X, Jiang Y, Tsou DA, Li S. Derivation of smooth muscle cells with neural crest origin from human induced pluripotent stem cells. Cells Tissues Organs. 2012; 195: 5–14. https://doi.org/10.1002/jcp.25437 PMID: 22005509

36. Kreitzer FR, Salomonis N, Sheehan A, Huang M, Park JS, Spindler MJ et al. A robust method to derive functional neural crest cells from human pluripotent stem cells. Am J Stem Cells. 2013; 2: 119–131. PMID: 23862100

37. Tomokiyo A, Hynes K, Ng J, Menicanin D, Camp E, Arthur A et al. Generation of Neural Crest-Like Cells From Human Periodontal Ligament Cell-Derived Induced Pluripotent Stem Cells. J Cell Physiol. 2017; 232: 402–416. https://doi.org/10.1002/jcp.25437 PMID: 27206577

38. Michael D, Wagoner MD, Bohrer LR, Aldrich BT, Greiner MA, Mullins RF et al. Feeder-free differentiation of cells exhibiting characteristics of corneal endothelium from human induced pluripotent stem cells. Biol Open. 2018; 7: 5. http://dx.doi.org/10.1242/bio.032102 PMID: 29685994

39. Bajpai R, Chen DA, Rada-Iglesias A, Zhang J, Xiong Y, Helms J et al. CHD7 cooperates with PBAF to control multipotent neural crest formation. Nature. 2010; 463: 958–962. https://doi.org/10.1038/nature08733 PMID: 20130577

40. Gallego RI, Pai AA, Tung J, Gilad Y. RNA-seq: impact of RNA degradation on transcript quantification. BMC Biol. 2014; 12: 42. https://dx.doi.org/10.1186%2F1741-7007-12-42 PMID: 24885439

41. Wang Z, Gerstein M, Snyder M. RNA-Seq: a revolutionary tool for transcriptomics. Nat Rev Genet. 2009; 10: 57–63. https://doi.org/10.1038/nrg2484 PMID: 19015660

42. Mortazavi A, Williams BA, McCue K, Schaeffer L, Wold B. Mapping and quantifying mammalian transcriptomes by RNA-Seq. Nat Methods. 2008; 5: 621–628. https://doi.org/10.1038/nmeth.1226 PMID: 18516045

43. Sauka-Spengler T, Meulemans D, Jones M, Bronner Fraser M. Ancient evolutionary origin of the neural crest gene regulatory network. Dev Cell. 2007; 13: 405–420. https://doi.org/10.1016/j.devcel.2007.08.005 PMID: 17765683

44. Nikitina N, Sauka-Spengler T, Bronner Fraser M. Dissecting early regulatory relationships in the lamprey neural crest gene network. Proc Natl Acad Sci U S A. 2008; 105: 20083 20088. https://dx.doi.org/10.1073%2Fpnas.0806009105 PMID: 19104059

45. Khudyakov J, Bronner Fraser M. Comprehensive spatiotemporal analysis of early chick neural crest network genes. Dev Dyn. 2009; 238: 716–723. https://doi.org/10.1002/dvdy.21881 PMID: 19235729

46. Hill RE, Jones PF, Rees AR, Sime CM, Justice MJ, Copeland NG et al. A new family of mouse homeo box-containing genes: molecular structure, chromosomal location, and developmental expression of Hox-7.1. Genes Dev. 1989; 3: 26–37. PMID: 2565278

47. Suzuki A, Ueno N, Hemmati Brivanlou A. Xenopus msx1 mediates epidermal induction and neural inhibition by BMP4. Development. 1997; 124: 3037–3044. PMID: 9272945

48. Simões-Costa M, McKeown SJ, Tan-Cabugao J, Sauka-Spengler T, Bronner ME. Dynamic and differential regulation of stem cell factor FoxD3 in the neural crest is Encrypted in the genome. PLoS Genet. 2012; 8: e1003142. https://doi.org/10.1371/journal.pgen.1003142 PMID: 23284303

49. Goulding MD, Chalepakis G, Deutsch U, Erselius JR, Gruss P. Pax-3, a novel murine DNA binding protein expressed during early neurogenesis. EMBO J. 1991; 10: 1135–1147. PMID: 202218

50. Bang AG, Papalopulu N, Goulding MD, Kintner C. Expression of Pax-3 in the lateral neural plate is dependent on a Wnt-mediated signal from posterior nonaxial mesoderm. Dev Biol. 1991; 212: 366–380. https://doi.org/10.1006/dbio.1999.9319 PMID: 10433827

51. Alkobtawi M, Ray H, Barriga EH, Moreno M, Kerney R, Monsoro-Burq AH et al. Characterization of Pax3 and Sox10 transgenic Xenopus laevis embryos as tools to study neural crest development. Dev Biol. 2018; 17: https://doi.org/10.1016/j.ydbio.2018.02.020 PMID: 29522707

52. Maczkowiak F, Matéos S, Wang E, Roche D, Harland R, Monsoro Burq AH. The Pax3 and Pax7 paralogs cooperate in neural and neural crest patterning using distinct molecular mechanisms, in Xenopus laevis embryos. Dev Biol. 2010; 340: 381–396. https://doi.org/10.1016/j.ydbio.2010.01.022 PMID: 20116373

53. Krishnakumar R, Chen AF, Pantovich MG, Danial M, Parchem RJ, Labosky PA et al. FOXD3 Regulates Pluripotent Stem Cell Potential by Simultaneously Initiating and Repressing Enhancer Activity. Cell Stem Cell. 2016; 18: 104–117. https://doi.org/10.1016/j.stem.2015.10.003 PMID: 26748757

54. Desanlis I, Felstead HL, Edwards DR, Wheeler GN. ADAMTS9, a member of the ADAMTS family, in Xenopus development. Gene Expr Patterns. 2018; 29: 72–81. https://doi.org/10.1016/j.gep.2018.06.001 PMID: 29935379

55. Porter S, Clark IM, Kevorkian L, Edwards DR. The ADAMTS metalloproteinases. Biochem J. 2005; 386: 15–27. https://doi.org/10.1042/BJ20040424 PMID: 15554875

56. Hubmacher D, Apte SS. ADAMTS proteins as modulators of microfibril formation and function. Matrix Biol. 2015; 47; 34–43. https://doi.org/10.1016/j.matbio.2015.05.004 PMID: 25957949

57. Chen WV, Maniatis T. Clustered protocadherins. Development. 2013; 140; 3297–302. https://doi.org/10.1242/dev.090621 PMID: 23900538

58. Okita K, Ichisaka T, Yamanaka S. Generation of germline competent induced pluripotent stem cells. Nature. 2007; 448; 313–317. https://doi.org/10.1038/nature05934 PMID: 17554338

59. Ishii M, Arias AC, Liu L, Chen YB, Bronner ME, Maxson RE. A stable cranial neural crest cell line from mouse. Stem Cells Dev. 2012; 21; 3069–3080. https://doi.org/10.1089/scd.2012.0155 PMID: 22889333

60. Antonellis A, Bennett WR, Menheniott TR, Prasad AB, Lee-Lin SQ; NISC Comparative Sequencing Program et al. Deletion of long-range sequences at Sox10 compromises developmental expression in a mouse model of Waardenburg-Shah (WS4) syndrome. Hum Mol Genet. 2006; 15; 259–271. https://doi.org/10.1093/hmg/ddi442 PMID: 16330480

61. Betancur P, Bronner-Fraser M, Sauka Spengler T. Genomic code for Sox10 activation reveals a key regulatory enhancer for cranial neural crest. Proc Natl Acad Sci U S A. 2010; 107; 3570–3575. https://doi.org/10.1073/pnas.0906596107 PMID: 20139305

62. Rinon A, Molchadsky A, Nathan E, Yovel G, Rotter V, Sarig R et al. p53 coordinates cranial neural crest cell growth and epithelial-mesenchymal transition/delamination processes. Development. 2011; 138: 1827–1838. https://doi.org/10.1242/dev.053645 PMID: 21447558

63. Hari L, Miescher I, Shakhova O, Suter U, Chin L, Taketo M et al. Temporal control of neural crest lineage generation by Wnt/β-catenin signaling. Development. 2012; 139: 2107–2117. https://doi.org/10.1242/dev.073064 PMID: 22573620

64. Murko C, Bronner ME. Tissue specific regulation of the chick Sox10E1 enhancer by different Sox family members. Dev Biol. 2016; 422: 47–57. https://doi.org/10.1016/j.ydbio.2016.12.004 PMID: 28012818

65. Spokony RF, Aoki Y, Saint-Germain N, Magner-Fink E, Saint Jeannet JP. The transcription factor Sox9 is required for cranial neural crest development in Xenopus. Development. 2002; 129: 421–432. PMID: 11807034

66. Perez-Alcala S, Nieto MA, Barbas JA. LSox5 regulates RhoB expression in the neural tube and promotes generation of the neural crest. Development. 2004; 131: 4455–4465. https://doi.org/10.1242/dev.01329 PMID: 15306568

67. Barembaum M, Bronner ME. Identification and dissection of a key enhancer mediating cranial neural crest specific expression of transcription factor, Ets-1. Dev Biol. 2013; 382:

68. Nagai T, Aruga J, Takada S, Günther T, Spörle R, Schughart K et al. The expression of the mouse Zic1, Zic2, and Zic3 gene suggests an essential role for Zic genes in body pattern formation. Dev Biol. 1997; 182: 299–313. https://doi.org/10.1006/dbio.1996.8449 PMID: 9070329

69. Teslaa JJ, Keller AN, Nyholm MK, Grinblat Y. Zebrafish Zic2a and Zic2b regulate neural crest and craniofacial development. Dev Biol. 2013; 380: 73–86. https://doi.org/10.1016/j.ydbio.2013.04.033 PMID: 23665173

70. Das A, Crump JG. Bmps and id2a act upstream of Twist1 to restrict ectomesenchyme potential of the cranial neural crest. PLoS Genet.2012; 8: e1002710. https://doi.org/10.1371/journal.pgen.1002710 PMID: 22589745

71. Machon O, Masek J, Machonova O, Krauss S, Kozmik Z. Meis2 is essential for cranial and cardiac neural crest development. BMC Dev Biol. 2015;15. https://doi.org/10.1186/s12861-015-0093-6 PMID: 26545946

72. Mitchell PJ, Timmons PM, Hébert JM, Rigby PW, Tjian R. Transcription factor AP-2 is expressed in neural crest cell lineages during mouse embryogenesis. Genes Dev. 1991; 5: 105–119. PMID: 1989904

73. Shen H, Wilke T, Ashique AM, Narvey M, Zerucha T, Savino E. Chicken transcription factor AP-2: cloning, expression and its role in outgrowth of facialprominences and limb buds. Dev Biol. 1997; 188: 248–266. https://doi.org/10.1006/dbio.1997.8617 PMID: 9268573

74. Luo T, Lee YH, Saint-Jeannet JP, Sargent TD. Induction of neural crest in Xenopus by transcription factor AP2alpha. Proc Natl Acad Sci U S A. 2003; 100: 532–537. https://doi.org/10.1073/pnas.0237226100 PMID: 12511599

75. de Crozé N, Maczkowiak F, Monsoro Burq AH. Reiterative AP2a activity controls sequential steps in the neural crest gene regulatory network. Proc Natl Acad Sci U S A. 2011; 108: 155–160. https://dx.doi.org/10.1073%2Fpnas.1010740107 PMID: 21169220

76. Wang WD, Melville DB, Montero-Balaguer M, Hatzopoulos AK, Knapik EW. Tfap2a and Foxd3 regulate early steps in the development of the neural crest progenitor population. Dev Biol. 2011; 360: 173–185. https://doi.org/10.1016/j.ydbio.2011.09.019 PMID: 21963426

77. Powell DR, Hernandez-Lagunas L, LaMonica K, Artinger KB. Prdm1a directly activates foxd3 and tfap2a during zebrafish neural crest specification. Development. 2013; 140: 3445–3455. https://doi.org/10.1242/dev.096164 PMID: 23900542

78. Yang L, Zhang H, Hu G, Wang H, Abate-Shen C, Shen MM. An early phase of embryonic Dlx5 expression defines the rostral boundary of the neural plate. J Neurosci. 1998; 18: 8322–8330. PMID: 9763476

79. Luo T, Matsuo-Takasaki M, Lim JH, Sargent TD. Differential regulation of Dlx gene expression by a BMP morphogenetic gradient. Int J Dev Biol. 2001; 45: 681–684. PMID:11461005

80. Li B, Kuriyama S, Moreno M, Mayor R. The posteriorizing gene Gbx2 is a direct target of Wnt signalling and the earliest factor in neural crest induction. Development. 2009; 136: 3267–3278. https://doi.org/10.1242/dev.036954 PMID: 19736322

81. Nakata K, Nagai T, Aruga J, Mikoshiba K. Xenopus Zic family and its role in neural crest development. Mech Dev. 1998; 75: 43–51. https://doi.org/10.1016/S0925-4773(98)00073 PMID: 9739105

82. Dottori M, Gross MK, Labosky P, Goulding M. The winged helix transcription factor Foxd3 suppresses interneuron differentiation and promotes neural crest cell fate. Development. 2001; 128: 4127–4138. PMID: 11684651

83. Kos R, Reedy MV, Johnson RL, Erickson CA. The winged-helix transcription factor FoxD3 is important for establishing the neural crest lineage and repressing melanogenesis in avian embryos. Development. 2001; 128: 1467–1479. PMID: 11262245

84. Wilson YM, Richards KL, Ford-Perriss ML, Panthier JJ, Murphy M. Neural crest cell lineage segregation in the mouse neural tube. Development. 2004; 131: 6153–6162. https://doi.org/10.1242/dev.01533 PMID: 15548576

85. Liu L, Chong SW, Balasubramaniyan NV, Korzh V, Ge R. Platelet-derived growth factor receptor alpha (pdgfr-α) gene in zebrafish embryonic development. Mech Dev. 2002; 116: 227–230. https://doi.org/10.1016/S0925-4773(02)00142-9 PMID: 12128230

86. Liu KJ, Harland RM. Cloning and characterization of Xenopus Id4 reveals differing roles for Id genes. Dev Biol. 2003; 264: 339–351. https://doi.org/10.1016/j.ydbio.2003.08.017 PMID: 14651922

87. Figueiredo AL, Maczkowiak F, Borday C, Pla P, Sittewelle M, Pegoraro C et al. PFKFB4 control of AKT signaling is essential for premigratory and migratory neural crest formation. Development. 2017; 144: 4183–4194. https://doi.org/10.1242/dev.157644 PMID: 29038306

88. Yang X, Li J, Zeng W, Li C, Mao B. Elongator Protein 3 (Elp3) stabilizes Snail1 and regulates neural crest migration in Xenopus. Sci Rep. 2016; 6: 26238. https://doi.org/10.1038/srep26238 PMID: 27189455

89. Sefton M, Sánchez S, Nieto MA. Conserved and divergent roles for members of the Snail family of transcription factors in the chick and mouse embryo. Development. 1998; 125: 3111–3121. PMID: 9671584

90. del Barrio MG, Nieto MA. Overexpression of Snail family members highlights their ability to promote chick neural crest formation. Development. 2002; 129: 1583–1593. PMID: 11923196

91. Aybar MJ, Nieto MA, Mayor R. Snail precedes Slug in the genetic cascade required for the specification and migration of the Xenopus neural crest. Development. 2003; 130: 483–494. http://doi.org/10.1242/dev.00238 PMID: 12490555

92. Nieto MA, Sargent MG, Wilkinson DG, Cooke J. Control of cell behavior during vertebrate development by Slug, a zinc finger gene. Science. 1994; 264: 835–859. http://doi.org/10.1126/science.7513443 PMID: 7513443

93. Jiang R, Lan Y, Norton CR, Sundberg JP, Gridley T. The Slug gene is not essential for mesoderm or neural crest development in mice. Dev Biol. 1998; 198: 277–285. https://doi.org/10.1016/S0012-1606(98)80005-5 PMID: 9659933

94. Tien CL, Jones A, Wang H, Gerigk M, Nozell S, Chang C. Snail2/Slug cooperates with Polycomb repressive complex 2 (PRC2) to regulate neural crest development. Development. 2015; 142: 722–731. https://doi.org/10.1242/dev.111997 PMID: 25617436

95. Martin BL, Harland PM. Hypaxial muscle migration during primary myogenesis in Xenopus laevis. Dev Biol. 2001; 239: 270–280. https://doi.org/10.1006/dbio.2001.0434 PMID: 11784034

96. Cheung M, Briscoe J. Neural crest development is regulated by the transcription factor Sox9. Development. 2003; 130: 5681–5693. https://doi.org/10.1242/dev.00808 PMID: 14522876

97. Cheung M, Chaboissier MC, Mynett A, Hirst E, Schedl A, Briscoe J. The transcriptional control of trunk neural crest induction, survival, and delamination. Dev Cell. 2005; 8: 179–192. https://doi.org/10.1016/j.devcel.2004.12.010 PMID: 15691760

98. Honoré SM, Aybar MJ, Mayor R. Sox10 is required for the early development of the prospective neural crest in Xenopus embryos. Dev Biol. 2003; 260: 79–96. https://doi.org/10.1016/S0012-1606(03)00247-1 PMID: 12885557

99. McKeown SJ, Lee VM, Bronner-Fraser M, Newgreen DF, Farlie PG. Sox10 overexpression induces neural crest-like cells from all dorsoventral levels of the neural tube but inhibits differentiation. Dev Dyn. 2005; 233: 430–444. https://doi.org/10.1002/dvdy.20341 PMID: 15768395

100. Prasad MK, Reed X, Gorkin DU, Cronin JC, McAdow AR, Chain K et al. SOX10 directly modulates ERBB3 transcription via an intronic neural crest enhancer. BMC Dev Biol. 2011; 11. https://doi.org/10.1186/1471-213X-11-40 PMID: 21672228

101. Baggiolini A, Varum S, Mateos JM, Bettosini D, John N, Bonalli M et al. Premigratory and migratory neural crest cells are multipotent in vivo. Cell Stem Cell. 2015; 16: 314–322. https://doi.org/10.1016/j.stem.2015.02.017 PMID: 25748934

102. McKinney MC, McLennan R, Kulesa PM. Angiopoietin 2 signaling plays a critical role in neural crest cell migration. BMC Biol. 2016; 14. https://doi.org/10.1186/s12915-016-0323-9 PMID: 27978830

103. Lee HO, Levorse JM, Shin MK. The endothelin receptor-B is required for the migration of neural crest-derived melanocyte and enteric neuron precursors. Dev Biol. 2003; 259: 162 175. https://doi.org/10.1016/S0012-1606(03)00160-X PMID: 12812796

104. Giovannone D, Ortega B, Reyes M, El-Ghali N, Rabadi M, Sao S et al. Chicken trunk neural crest migration visualized with HNK1. Acta Histochem. 2015; 117: 255–266. https://doi.org/10.1016/j.acthis.2015.03.002 PMID: 25805416

105. Zuhdi N, Ortega B, Giovannone D, Ra H, Reyes M, Asención V et al. Slits Affect the Timely Migration of Neural Crest Cells Via Robo Receptor. Dev Dyn. 2012; 241: 1274 1288. https://dx.doi.org/10.1002%2Fdvdy.23817 PMID: 22689303

106. Chiovaro F, Chiquet-Ehrismann R, Chiquet M. Transcriptional regulation of tenascin genes. Cell Adh Migr. 2015; 9: 34–47. https://dx.doi.org/10.1080%2F19336918.2015.1008333 PMID: 25793574

107. Taneyhill AL, Coles EG, Bronner Fraser M. Snail2 directly represses cadherin6B during epithelial-to-mesenchymal transitions of the neural crest. Development. 2007; 134: 1480–1490. https://dx.doi.org/10.1242%2Fdev.02834 PMID: 17344227

108. Groysman M, Shoval I, Kalcheim C. A negative modulatory role for rho and rho-associated kinase signaling in delamination of neural crest cells. Neural Develop. 2008; 3: 27. https://dx.doi.org/10.1186%2F1749-8104-3-27 PMID: 18945340

109. Vega FM, Thomas M, Reymond N, Ridley AJ. The Rho GTPase RhoB regulates cadherin expression and epithelial cell-cell interaction. Cell Commun Signal. 2015; 13: 6. https://doi.org/10.1186/s12964-015-0085-y PMID: 25630770

110. Liu Q, Dalman MR, Sarmah S, Chen S, Chen Y, Hurlbut AK et al. Cell adhesion molecule cadherin-6 function in zebrafish cranial and lateral line ganglia development. Dev Dyn. 2011; 240: 1716–26. https://doi.org/10.1002/dvdy.22665 PMID: 21584906

111. Chiovaro F, Chiquet-Ehrismann R, Chiquet M. Transcriptional regulation of tenascin genes. Cell Adh Migr. 2015; 9: 34–47. https://dx.doi.org/10.1080%2F19336918.2015.1008333 PMID: 25793574

112. Tomczuk M, Takahashi Y, Huang J, Murase S, Mistretta M, Klaffky E et al. Role of multiple beta1 integrins in cell adhesion to the disintegrin domains of ADAMs 2 and 3. Exp Cell Res. 2003; 290: 68–81. https://doi.org/10.1016/S0014-4827(03)00307-0 PMID: 14516789

113. Dennis AR, McLennan R, Jessica MT, Craig LS, Jeffrey SH, Kulesa PM. The neural crest cell cycle is related to phases of migration in the head. Development. 2014; 141: 1095–1103. https://dx.doi.org/10.1242%2Fdev.098855 PMID: 24550117

